# Extending site-based observations to predict the spatial patterns of vegetation structure and composition

**DOI:** 10.1101/715797

**Authors:** Megan J McNellie, Ian Oliver, Simon Ferrier, Graeme Newell, Glenn Manion, Peter Griffioen, Matt White, Terry Koen, Michael Somerville, Philip Gibbons

## Abstract

**Context:** Conservation planning and land management are inherently spatial processes that are most effective when implemented over large areas.

**Objectives:** Our objectives were to (i) use existing plot data to aggregate species inventories to growth forms and derive indicators of vegetation structure and composition and ii) generate spatially-explicit, continuous, landscape scaled models of these discrete vegetation indicators, accompanied by maps of model uncertainty.

**Method:** Using a case study from New South Wales, Australia, we aggregated floristic observations from 7234 sites into growth forms. We trained ensembles of artificial neural networks (ANN) to predict the distribution of these indicators over a broad region covering 11.5 million hectares. Importantly, we show spatially explicit models of uncertainty so that end-users have a tangible and transparent means of assessing models.

**Results:** Our key findings were firstly, widely available site-based floristic records can be used to derive aggregated indicators of the structure and composition of plant growth forms. Secondly, ANNs are a powerful method to predict continuous patterns in complex, non-linear data (Pearson’s correlation coefficient 0.83 (total native vegetation cover) to 0.42 (forb cover)). Thirdly, maps of the standardised residual error give insight into model performance and provide an assessment of model uncertainty in specific locations.

**Conclusions:** Spatially explicit, continuous representations of vegetation composition and structural complexity can add considerable value to conventional maps of vegetation extent or community type. This application has the potential to enhance the capacity for conservation planners, landscape managers and policy-makers to make informed decisions across landscape and regional scales.

## Introduction

Conservation planning and land management are inherently spatial processes, and to be effective, they need to be implemented over extensive geographic regions. Vegetation extent and community type are often the primary spatial information used to underpin decision-making at broad scales. Typically, maps of vegetation extent depict binary classifications such as ‘extant versus cleared’; ‘woody versus non-woody’ or ‘native versus exotic’; and maps of vegetation communities depict discrete boundaries of community types, each deemed to be internally homogeneous in terms of floristic composition, and distinct from other types.

Maps of extent and type have been used to assess the impacts of land use (e.g. Hansen et al. 2013), model the distribution of species (e.g. Ferrier et al. 2002), inform systematic conservation planning (e.g. Margules and Pressey 2000), implement conservation prioritization (e.g. Moilanen et al. 2011; Myers et al. 2000), assess wildfire risk (e.g. McKenzie et al. 2004) and estimate standing carbon stocks (e.g. Chave et al. 2005; Scurlock and Hall 1998). For example, in their work identifying global hotspots, Myers et al. (2000) identified the extent of remaining vegetation types as one of the determinants of hotspot status. Myers et al. (2000) and many others, pragmatically defer to such maps because vegetation extent and community type are often the only spatially explicit datasets available across large extents.

However, maps with delineated boundaries imply vegetation is discontinuous (De Cáceres and Wiser 2011). Binary or categorical maps seldom represent the continuous characteristics of vegetation, such as foliage cover and species richness. Nor do they directly convey information on total native plant cover or total native abundance, or on the multi-layered components of plant structure or composition of discrete growth forms. Often the discrete structural elements, or the relative abundance and composition, of different plants are difficult to discern from maps of vegetation communities.

The structural complexity and composition of vegetation is a function of climate and soils, geographic position in the landscape, biogeographic processes and of past disturbance (both human-induced and natural stochastic events) (Specht and Specht 2002). Structural complexity and heterogeneity are associated with faunal diversity and distribution (Lindenmayer et al. 2000; McElhinny et al. 2006a; Tews et al. 2004); can be used to judge the risk of wildfire by assessing co-occurring plant growth forms (Pyne et al. 1996) and can be used to account for the above-ground biomass that contributes to carbon stocks (Scurlock and Hall 1998). The compositional elements of vegetation at a site, such as native species richness underlie conservation strategies (Fleishman et al. 2006) and can be used to identify threats to biodiversity (Evans et al. 2011; Hooper et al. 2005).

Growth forms (such as trees, shrubs, grasses or forbs) represent aggregations of species based on plant form and function and provide a useful approach for representing the multi-layer structural complexity and the relative abundance and composition of vegetation. Globally, site-based floristic inventories that record individually observed plant species (Bruelheide et al. 2019; Franklin et al. 2017) and their growth form allocations (Engemann et al. 2016; Oliver et al. 2019) are becoming easier to access.

The first objective of our study was to use existing floristic inventories recorded from vegetation plots and synthesise these lists of species into indicators of growth form structure and composition. The second objective was use predictive models to interpolate the ecological patterns and map these predictions at a landscape scale. We coupled aggregated observational data with predictor surfaces that influence the growth and morphology of vegetation, as well as predictor surfaces that reflect landscape-scaled disturbance. Of the range of modelling methods available to interpolate site-based observation data (Elith et al. 2006), we used ensembles of artificial neural network (ANN) models. ANNs are advantageous for ecological applications where data do not meet parametric statistical assumptions and the relationships between the response data and the predictor surfaces are complex, unknown or non-linear (Bishop 1995; Fielding 1999).

These resultant predictive maps of individual vegetation indicators represent spatially-explicit, continuous characteristics of vegetation structure and composition. Furthermore, we mapped the spatial distribution of the standardised residual errors to show explicitly where our models over- or under-predict. This is a key asset in ensuring that the strengths and limitations of these spatially-explicit predictions are clear to end-users. This information can support a broad range of landscape scaled applications and help conservation planners and landscape managers make better informed decisions.

## Methods

### Study area

Our study area of 115 000 km^2^ is located in north-eastern New South Wales (NSW), Australia (Figure 1). The area is characterised by privately-owned land used for agriculture (52% land used for grazing and 23% used for cultivation, including irrigated cropping). Extractive industries (including coal seam gas exploration and open-cut coal mining) are emerging land uses. Less than 10% of the area is protected within National Park or public reserve. The biophysical landscape includes elevations up to 1510 m in the eastern ranges to semi-arid regions on the western plains. Vegetation formations are diverse, ranging from closed rainforest in the eastern, elevated regions to arid shrublands and grasslands on the drier, western plains (Keith 2004).

**Fig 1.**
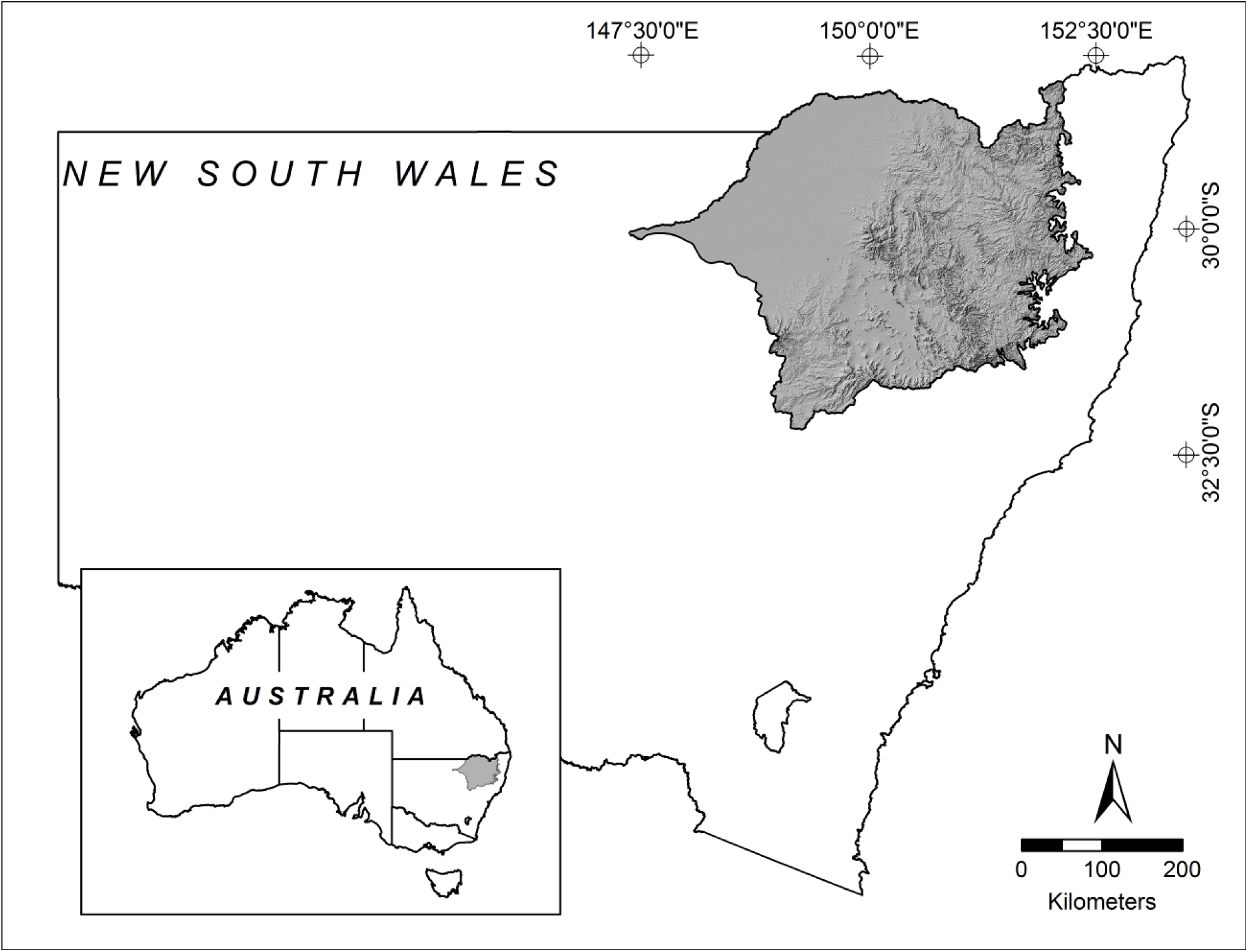
Location of the study area showing topographic relief - northern New South Wales, Australia.

Figure 2 outlines the five components of this study: (i) transforming site-based data into vegetation indicators; (ii) sourcing spatial layers that represent environmental and disturbance gradients; (iii) training ANN models; (iv) using the trained ANN model to predict indicators across the whole landscape; and (v) rendering the average results from ensembles of predicted models into a spatially explicit map for each indicator, accompanied by a mapped estimate of the standardised residual error.

**Fig 2.**
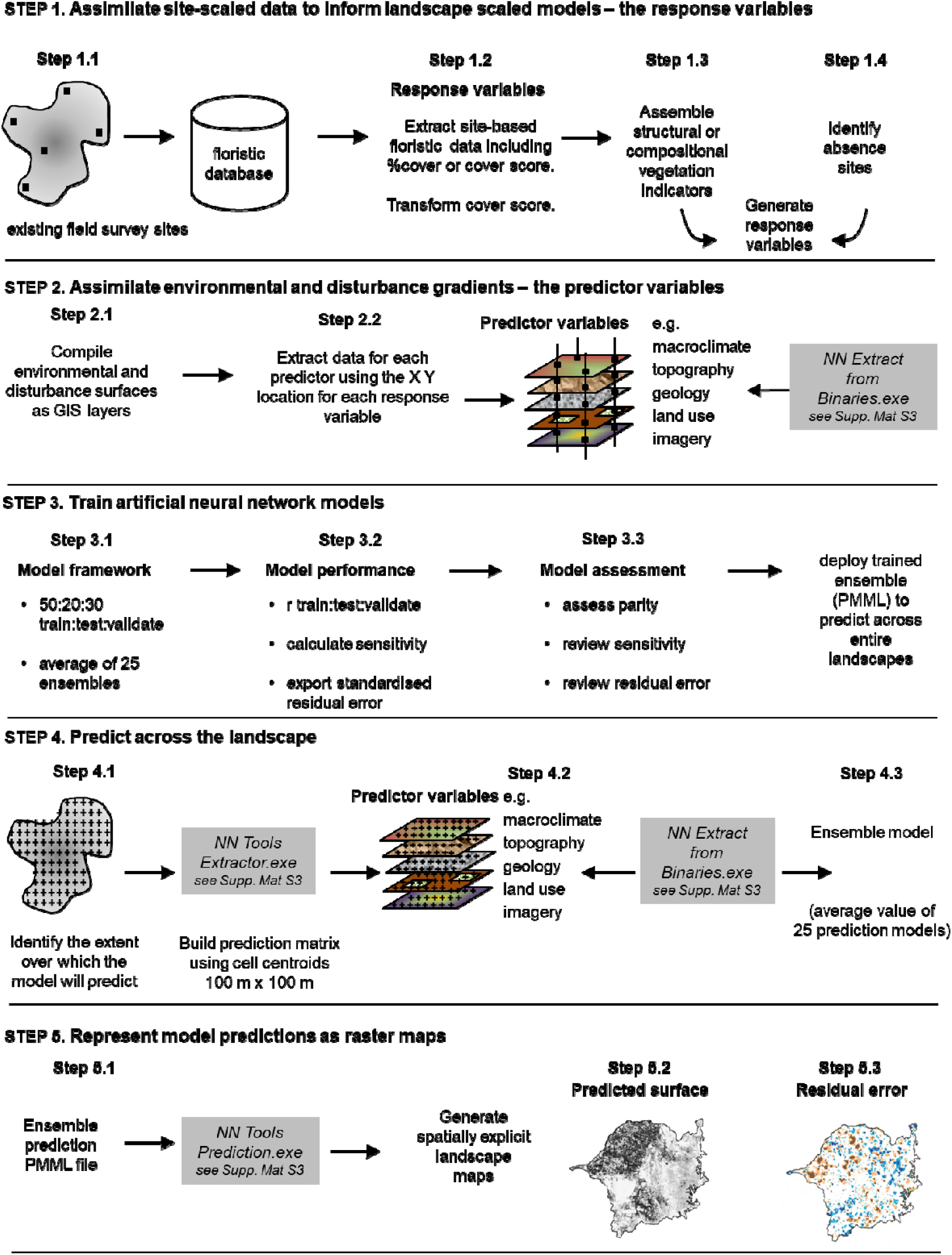
Workflow showing an overview of the five components required to transform floristic data to deliver landscape-scaled models of vegetation indicators

### Site-based data to inform landscape-scaled models – the response variables

We extracted floristic records from in the BioNet Database (http://www.bionet.nsw.gov.au). This database contains over 100 000 geo-referenced, site-based surveys of vascular plants across the state of NSW. These floristic data have been collected for a multitude of reasons, mostly to inventory, circumscribe and map vegetation communities. We extracted only floristic records that were surveyed from a fixed area (0.04 ha); included either a percentage foliage cover estimate (<1%–100%) or a cover-abundance score (e.g. 1–6 Braun-Blanquet cover abundance score) for each plant species and contained no missing information (Step 1.1 in Figure 2). The suite of sites extracted from the database spanned a survey period from 1986 to 2011.

Floristic records were checked (e.g. metadata were reviewed to identify only records that were collected systematically and recorded full floristic data) and cleaned (e.g. sites with missing data were removed). Floristic data with Braun-Blanquet cover-abundance scores were converted to percentage cover values following methods described in McNellie et al. (2019) (Step 1.2 in Figure 2). All species were assigned as either native or exotic. We accessed an existing framework for allocating native species to one of six major growth form groups (trees, shrubs, grasses and grass-like, forbs, ferns, and remaining ‘others’ (Oliver et al. 2019) (Step 1.3 in Figure 2). Here we have excluded ferns and other growth form groups because there were too few observations and we focused on four dominant growth form groups: trees, shrubs, grass and grass-like (hereafter referred to as grasses) and forbs.

Structural indicators were calculated by summing the foliage cover estimates across all native species. Compositional indicators were calculated as the count of all native species (after aggregating subspecies and varieties). Outliers were identified by plotting the distribution data. Sites with abnormally high summed cover (most likely due to human error in making visual estimates of cover) were identified and excluded from cover models. In total, ten vegetation indicators were generated: total native cover; total native richness; as well as cover and richness of trees, shrubs, grasses and forbs.

To improve predictive accuracy and to build a representative dataset, we created an additional set of absence sites (Step 1.4 in Figure 2). Given our exclusive interest in the structural and compositional properties and characteristics of the native vegetation, our ‘absences’ were locations where there is virtually no perennial cover of native vegetation. Absence sites were important in this study as the training data were almost exclusively obtained from natural or near-natural vegetation and the objective of this study was to extrapolate our models across all land tenures, including both near-natural systems and small remnants of vegetation. This complementary set of absences in the response data were generated using on-screen digitising point registration using satellite imagery (2009 SPOT 5) as a backdrop to identify under-sampled land uses that contain no native terrestrial vegetation such as irrigation bays, water bodies, roads and infrastructure. Each absence site was attributed as having 0% native species cover and 0 native species richness.

### Environmental and disturbance surfaces – the predictor variables

We selected a suite of predictor surfaces that reflect: (a) environmental attributes that directly (temperature, moisture, radiation) or indirectly (geology, soils, topography) influence the resources and conditions controlling growth and morphology of vegetation (Austin 2002; Box 1981; Franklin 2009; Pressey et al. 2000) and (b) an *a priori* assessment of disturbance variables that modify and fragment vegetation (Step 2.1 in Figure 2 and Supplementary Material S1) (Foley et al. 2005).

#### i) Abiotic environmental surfaces

Climatic and topographic surfaces were derived from the Australian 1 second, smoothed digital elevation model (DEM-S) (Gallant et al. 2011). Raster surfaces were resampled to 25 m resolution to match the observational scale of the response data (Williams et al. 2012). Climatic variables were calculated using the MTHCLIM module in ANUCLIM v6.1 (Xu and Hutchinson 2013) for the 1921-1995 epoch. Detailed and additional information on predictor surfaces is provided in Supplementary Material S1.

#### ii) Contemporary disturbance surfaces

We used land use mapping as a surrogate for disturbance (Foley et al. 2005). We assigned mapped land use classes to seven major groups: grazing; cropping and horticulture; conservation areas; tree and shrub cover; remaining other land uses (which included all urban areas, roads, mining, power generation and areas used for intensive animal production); and rivers and wetlands. We also prepared continuous land use variables by calculating the distance (m) to the nearest land use or land modification (cropping, irrigation, grazing, mining or urban land uses). We compared model performance using either categorical or continuous land use variables.

Predictor variables derived from Landsat TM imagery were used only to inform models of total native vegetation cover and vegetation cover by growth form. Imagery from 2005 to 2012 was used to calculate normalized difference vegetation index (NDVI), foliage projected cover (FPC) and individual components of fractional groundcover (bare ground, green groundcover and brown groundcover) derived from multitemporal Landsat TM images. Consequently, the response data used to build and train models of total native vegetation cover and vegetation cover by growth form were a subset of the data where sites were surveyed between 2005 and 2012.

### Training the artificial neural network

The training matrix (Step 2.2 in Figure 2) was created by extracting the value for each predictor surface for each site with response data, as well as every absence site (see Supplementary Material S3 - NN Extract from Binaries.exe for software designed to build matrices). Lek et al. (1996), Olden et al. (2008), Özesmi et al. (2006) and Zhang et al. (1998) provide examples of working with ANN models within an ecological context and Bishop (1995), Fielding (1999) and Haykin (2009) provide comprehensive and authoritative texts on the principles of neural networks. The training matrix, absence sites and extracted values for all predictor surfaces are stored at <u>DOI://10.6084/m9.figshare.7730726</u>.

For each vegetation indicator, the response data were randomly split into three subsets: 50% of the data were used to train the model; 20% were used to test the model (by comparing the sum of squares error in the training and testing subsets). The remaining 30% were withheld (from training and testing) and used to independently validate the model (Step 3.1 in Figure 2). This form of data partitioning is a rigorous method for validating models (Fielding 1999). Additional information on the model architecture is detailed in Supplementary Material S2.

We trained artificial neural network models using 100 iterations whereby each iteration represented a different model architecture (e.g. number of hidden nodes). Each iteration of the model learns patterns or relationships between the response data and the predictor data (Bishop 1995). Of the 100 model iterations, the 25 highest performing models were combined and averaged to produce an ensemble model. The predictive performance of each ensemble model was evaluated by calculating the average Pearson’s correlation coefficient (*r*) for each of the training, testing and validation subsets (Step 3.2 in Figure 2). It is important to note that model performance was judged by determining how well the model performs when applied to new data (the validation subset). Parity between the correlation coefficients for the training and validation subsets gives an indication of how well the model has been trained (Step 3.3 in Figure 2). Sum of squares error was evaluated for each network model. Sensitivity analyses were reviewed to give an indication of the contribution of each predictor surface to the model (Step 3.3 in Figure 2). Sensitivity analyses are unit-less measurements and show how the model performs when each predictor is removed from analysis. Standardised residual error for each of the response data were reviewed to assess model uncertainty (see Step 5.2 in Figure 2). The standardised residual error can be interpreted like z-scores whereby values +/− 3 are regarded as outliers (Shekhar et al. 2003).

### Predicting the spatial patterns and relationships across the whole landscape

In this stage, we used raster analyses to treat every 100m grid cell in the study area (approximately 11.5 million grid cells) as a new, unknown, or unsurveyed site (Step 4.1 in Figure 2). The centroid of each grid cell was extracted using custom software (see Supplementary Material S3 - NN Extractor.exe) and the underlying values of the predictor surfaces were extracted to build the prediction matrix (see Supplementary Material S3 - NN Extract from Binaries.exe for details of the software designed to build this matrix). This matrix (sites x predictor variables) formed the ‘predict later’ (Ferrier and Guisan 2006) input table used to predict the trends and patterns learned from the training data site (Step 4.2 in Figure 2).

To build predictive models of each of the ten vegetation indicators, that is, to interpolate the spatial patterns and relationships across the whole landscape, each of the 25 training models per indicator (representing 25 trained networks) in Predictive Model Markup Language (PMML) file format were deployed to every grid cell in the prediction matrix which represented the new, unknown cases. These analyses produced 25 prediction models. The final predicted output for each grid cell was averaged to create a single ensemble model for each vegetation indicator (Step 4.3 in Figure 2).

### Representing the model predictions as maps

Predictions from the models were stored as PMML files. Custom software (Step 5.1 in Figure 2) (see Supplementary Material S3 - NN Tools Prediction.exe for software designed to build spatially explicit predictive models), transformed the PMML predicted model output into the format needed to create a raster map. Each vegetation indicator was mapped using the location (easting and northing) of each 100m x 100m cell centre (X, Y) and the predicted value (Z) (Step 5.2 in Figure 2). An overlay of non-native vegetation was used to mask out areas that have been identified as cleared (NSW Office of Environment and Heritage 2017).

To investigate the spatial patterns and behaviour of model predictions over the whole of landscape, we calculated the inverse distance weighting (IDW) (Shekhar et al. 2003) of the standardised residual to represent under- and over-prediction ‘error’ as a raster map (Step 5.3 in Figure 2). Negative standardised residual values indicate over-prediction and positive values indicate under-prediction. The spatial representation of the standardised residual error guides end-users where the modelled predictions have higher uncertainty.

## Results

### Response variables

The dataset used to train the models contained 7234 sites and an additional 3904 absence sites. The subset of sites that matched the temporal period of the remote sensing variables (and that were used to predict cover) was 3490 floristic sites and we selected a random subset of 1313 absence sites. Of the 3490 sites used as response data in the cover models, a further nine sites were excluded from the total native cover model because they were identified as outliers (>235% summed cover).

### Model assessment

The correlation coefficients, assessing the strength of correlation between predicted and observed values for of the training, testing and validation subsets for each modelled vegetation indicator are shown in Table 1. The correlation coefficient (*r*) for the validation subsets ranged from 0.83 (total native cover) to 0.42 (for forb cover). Parity between the correlation coefficients of the training subset and validation subsets show that models have been well-trained and performed well when applied to new data (the validation subset). The difference between the training and validation correlation coefficients was greater for cover models, with this difference ranging from 0.01 (forb cover) to 0.07 (shrub cover). The correlation coefficients for training and validation subsets in the richness models were much closer and the difference ranged from 0 (total native richness) to 0.02 (shrub and grass richness).

**Table 1:**
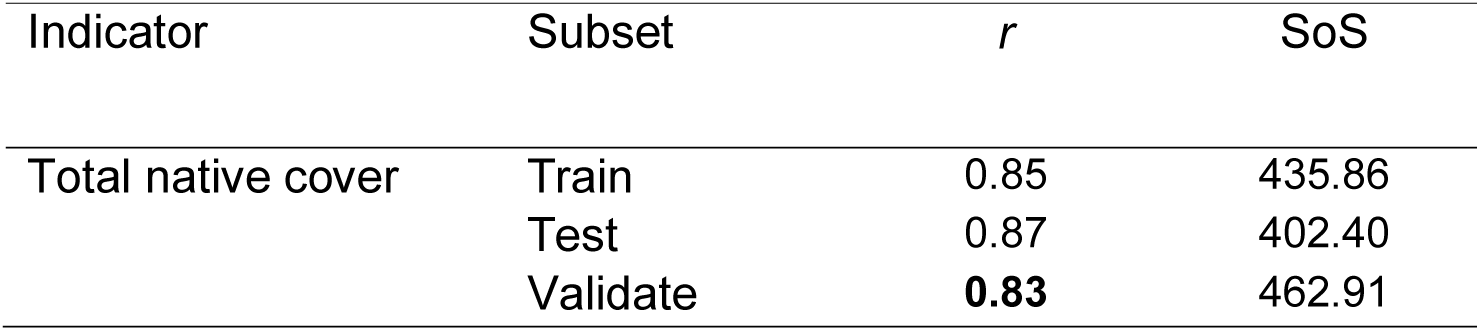
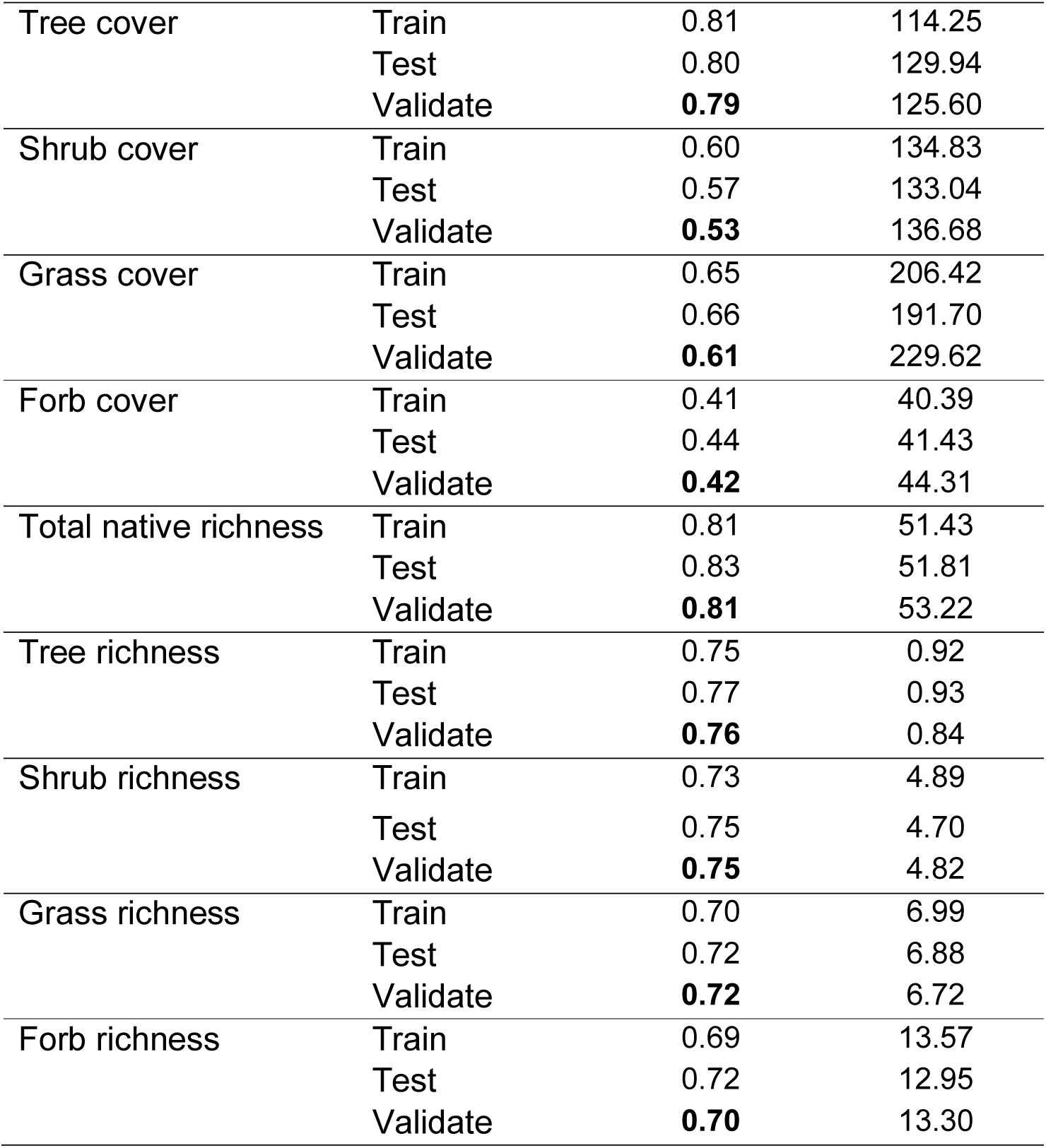
Averaged Pearson’s correlation coefficient (*r*) and sum of squares error (SoS) for ensemble models for each of the ten vegetation indicators. Correlation coefficients for the validation subset is shown in bold to highlight the validation results. Total number of observations (n) for total native cover model n = 3481 (with training subset n = 1741; test subset n = 696 and validation subset n = 1044); all other cover models n = 3490 (with training subset n = 1745; test subset n = 698 and validation subset n = 1047) and total number of observations for richness models n = 11132 (with training subset n = 5567; test subset n = 2226 and validation subset n = 3339).

| Indicator | Subset | $r$ | SoS |
| --- | --- | --- | --- |
| Total native cover | Train | 0.85 | 435.86 |
|  | Test | 0.87 | 402.40 |
|  | Validate | **0.83** | 462.91 |
| Tree cover | Train | 0.81 | 114.25 |
|  | Test | 0.80 | 129.94 |
|  | Validate | **0.79** | 125.60 |
| Shrub cover | Train | 0.60 | 134.83 |
|  | Test | 0.57 | 133.04 |
|  | Validate | **0.53** | 136.68 |
| Grass cover | Train | 0.65 | 206.42 |
|  | Test | 0.66 | 191.70 |
|  | Validate | **0.61** | 229.62 |
| Forb cover | Train | 0.41 | 40.39 |
|  | Test | 0.44 | 41.43 |
|  | Validate | **0.42** | 44.31 |
| Total native richness | Train | 0.81 | 51.43 |
|  | Test | 0.83 | 51.81 |
|  | Validate | **0.81** | 53.22 |
| Tree richness | Train | 0.75 | 0.92 |
|  | Test | 0.77 | 0.93 |
|  | Validate | **0.76** | 0.84 |
| Shrub richness | Train | 0.73 | 4.89 |
|  | Test | 0.75 | 4.70 |
|  | Validate | **0.75** | 4.82 |
| Grass richness | Train | 0.70 | 6.99 |
|  | Test | 0.72 | 6.88 |
|  | Validate | **0.72** | 6.72 |
| Forb richness | Train | 0.69 | 13.57 |
|  | Test | 0.72 | 12.95 |
|  | Validate | **0.70** | 13.30 |

### Predictor variables and sensitivity analysis

We evaluated treating land use as either a single categorical predictor or several individual continuous predictors. Our results (Supplementary Material S4) showed that correlation coefficients were higher and sum of squares errors were lower when categorical land use was used as a predictor to inform vegetation structure and composition. Sensitivity analysis showed that the categorical predictors (land use and great soil group) made the two highest contributions to training models. No predictors had a sensitivity of less than one (Table 2) indicating that there were no redundant predictor variables. Overall, the suite of richness models yielded higher sensitivity values for land use and soils, followed by climatic variables (mean minimum winter temperature and isothermality) and soil properties (percentage of clay or silt). Not surprisingly, remotely sensed predictor variables (foliage projective cover and normalised difference of vegetation index) had higher sensitivity values for the suite of cover models, especially total native cover and tree cover. However, for some of the cover attributes, such as forb cover, there were few strong predictor variables.

**Table 2:**
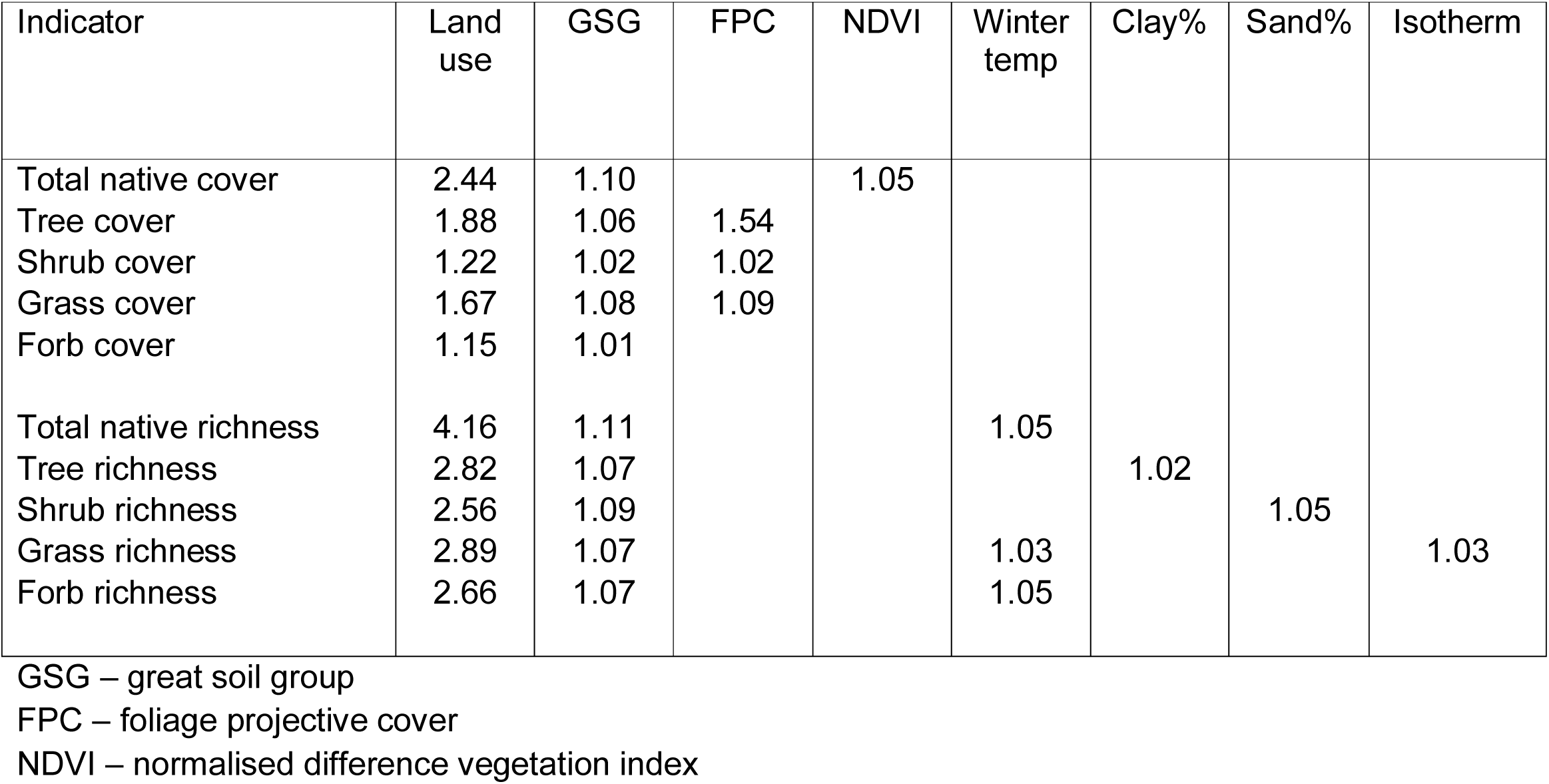
Sensitivity analysis showing the sensitivity averaged over 25 ensembles of ANN models for ten vegetation indicators. Only the highest three sensitivity values for each vegetation indicator are shown.

Interpolating spatial patterns and relationships across the whole landscape We created spatially explicit model-based predictions for ten vegetation indicators over the 11.5 million grid cells (100 m x 100 m) in the study area. The data file with location (X, Y) and the predicted value (Z) was displayed as a continuous surface raster map. Here we show an example (Figure 3) of the detailed (1:1 000 000 scale) prediction surface for total native richness (Figure 3a) and the associated estimate of standardised residual error in the prediction (Figure 3b), for a section of the study area (Figure 3c). Residual errors were classified to depict classes of under- and over-estimation. To further demonstrate how indicators of individual vegetation components can be modelled, Figure 4 shows a detailed example for shrub richness (1:1 000 000 scale) over the same area which illustrates the contribution of richness within this growth form to the overall richness. The spatially explicit maps for all ten vegetation indicators and their accompanying residual error maps are supplied as Supplementary Material S5.

**Fig 3.**
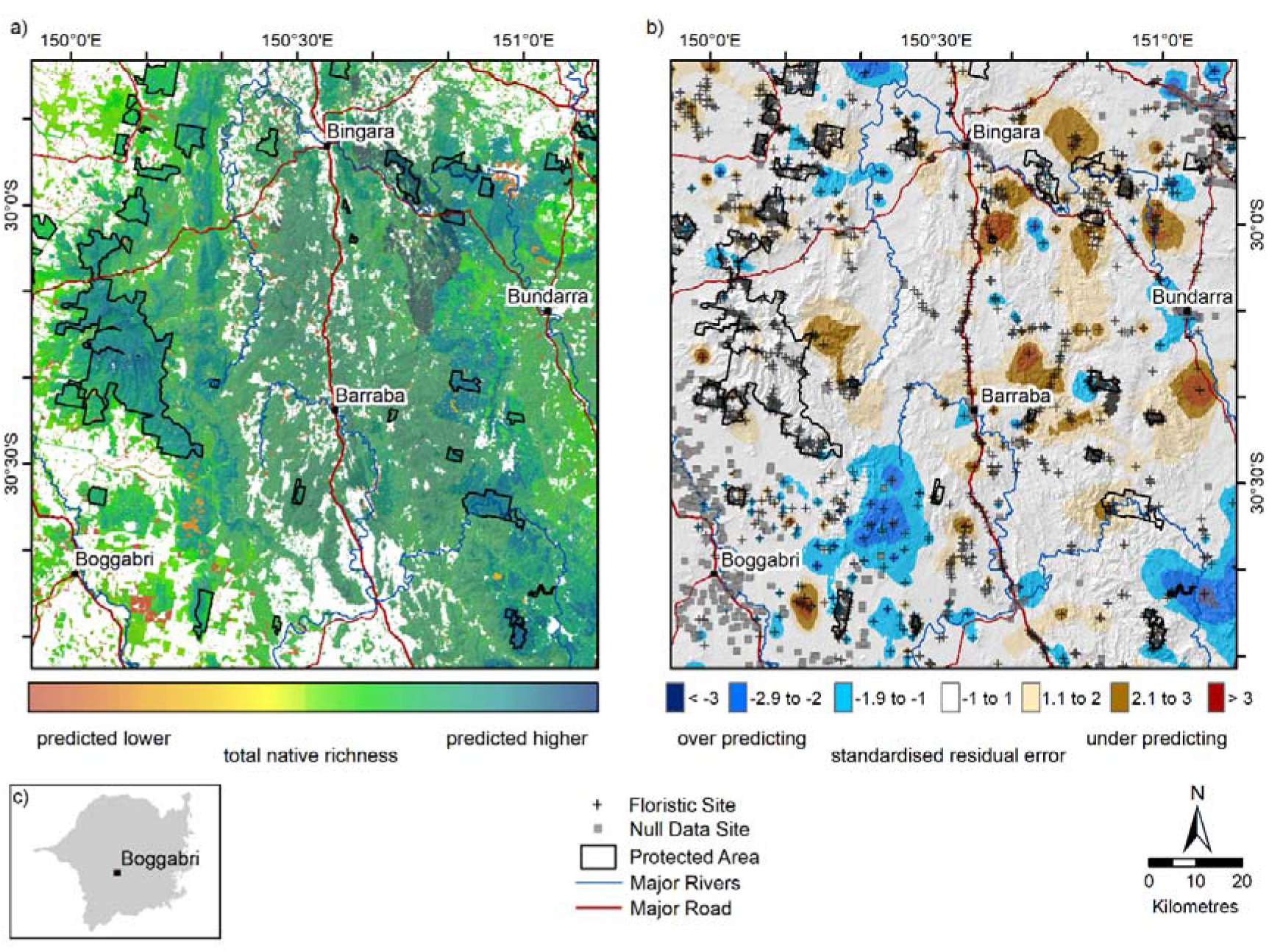
Fine-scale representation of predicted total native species richness showing the detail of the a) continuous surface for native species richness (count of all native species), b) standardised residual error for native species richness and c) a location diagram within the study area

**Fig 4.**
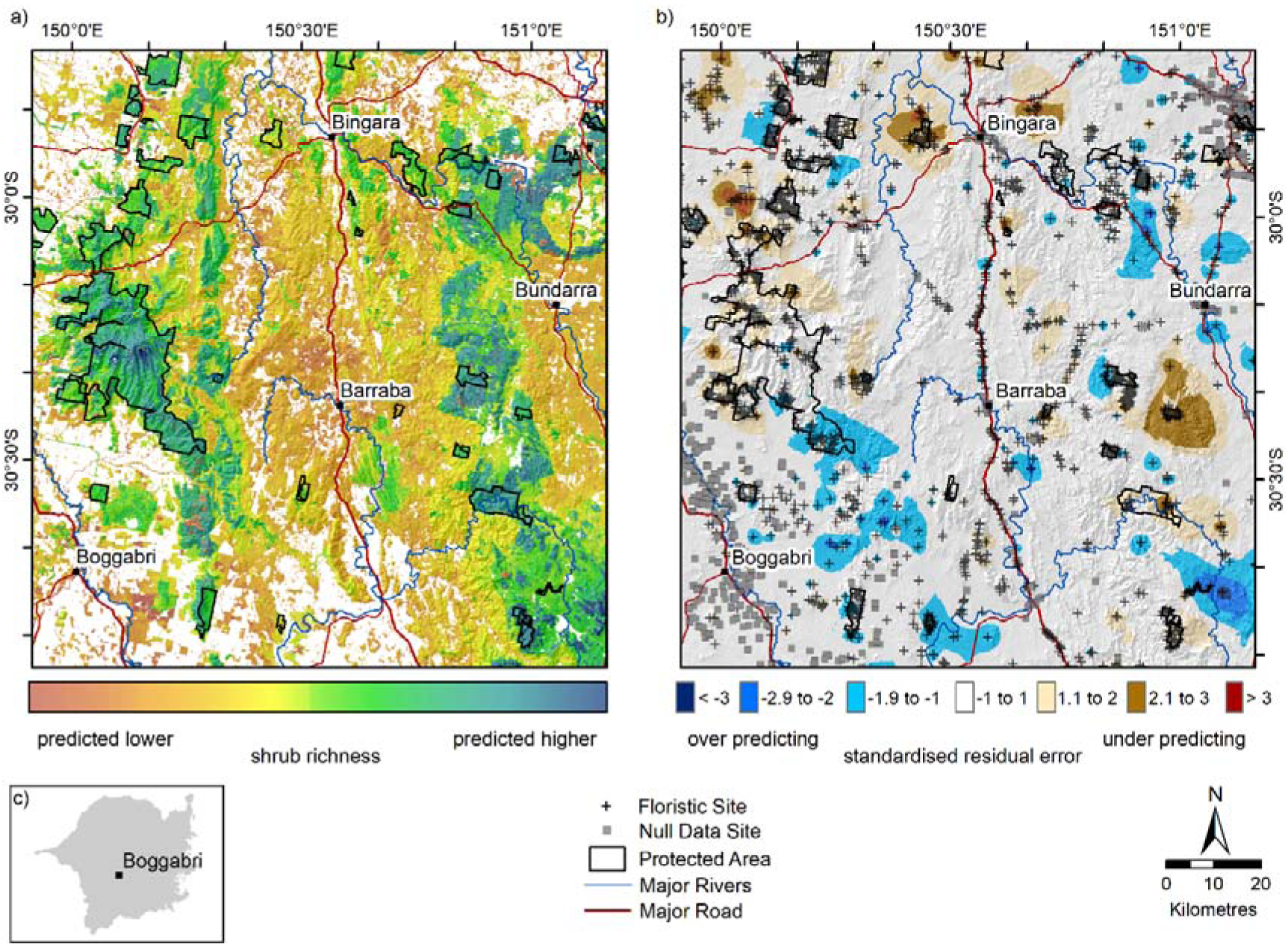
Fine-scale representation of predicted shrub richness showing the detail of the a) continuous surface for shrub richness, b) standardised residual error for shrub richness and c) a location diagram within the study area

## Discussion

### Extending site-data for predictive ecological modelling

Here we have shown how existing inventories of floristic data can be assembled into growth forms which can facilitate the modelling and more detailed mapping of different facets of vegetation structure and composition. The ever-growing volume of floristics site data (Bruelheide et al. 2019; Dengler et al. 2011; Peet et al. 2013; Schaminée et al. 2011) and large-scale synthesises of species to a growth form (Engemann et al. 2016; Oliver et al. 2019) has opened opportunities to explore how site-based observations can be extended to inform ecological models. Currently, much effort and attention has been applied to using these floristic data to inventory, describe and map vegetation community types at broad scales (e.g. Chytrý et al. 2011; Grossman et al. 1998; Mucina and van der Maarel 1989; Wiser et al. 2011), and these maps have been highly influential for supporting whole of landscape decision making (e.g. Myers et al. 2000).

However, approaches that aim to delineate boundaries around uniform types are subject to multiple sources of variation. Hearn et al. (2011) found that most errors in mapping vegetation boundaries were observed where neighbouring vegetation types had similar structure and composition. Delineating communities on a map is often an arbitrary decision that requires a degree of expert interpretation because most vegetation types intergrade across ecotones. Furthermore, Hearn et al. (2011) also found that experts differed in their opinions about which vegetation types were contained within the mapped boundaries, especially when vegetation structure and composition were similar (such as in shrub-dominated heaths or grasslands). Our spatially explicit representation of structure and composition of individual plant growth forms can overcome some of the sources of variation found in maps of vegetation communities. This alternative approach is unimpeded by bounded vegetation categories because we represent discrete growth forms as a continuous surface that can express the heterogeneity in vegetation structure and composition across the landscape.

### Evaluating the predictor data

Land use was a strong predictor for all vegetation structure and composition indices. Land use is a key driver underpinning the modification of natural landscapes (Fischer and Lindenmayer 2007; Foley et al. 2005; Newbold et al. 2016). Interestingly, when we investigated both categorical and continuous methods to represent land use data we found the categorical land use variable improved the correlation coefficients, reduced residual errors, made the greatest contribution to sensitivity analyses and resulted in realistic spatial representations of all indices across the landscape when compared to continuous representations of distance to different land use types. Lindegarth and Gamfeldt (2005) also found analyses using categorical variables provided an indication as to which factors were important.

This study addresses the challenge of selecting an appropriate and comprehensive set of predictor surfaces (Williams et al. 2012) by using predictors that are known to relate to growth and morphology of vegetation, such as temperature, light, topography, hydrology (Austin 1998; Box 1981), as well as predictors that are known to modify and fragment landscapes. The approach described here has the capacity to be broadened. Globally, compilations of gridded environmental surfaces that represent climate, soils or topography are readily available (e.g. for climatic data see WorldClim (Fick and Hijmans 2017); for soil data see SoilGrids (Hengl et al. 2017) or digital elevation models from which topographic predictors can be derived (e.g. ASTER Global Digital Elevation Model https://lpdaac.usgs.gov).

Despite our careful selection of predictor surfaces, there are limitations in using surrogates, such as land use, to represent disturbance. Firstly, for some land use categories, the extent to which native vegetation is modified is not uniform. For example, grazing by either native animals or livestock varies in its intensity and impact on native vegetation (Lunt et al. 2007; Olff and Ritchie 1998; Pausas and Bond 2019; Speed and Austrheim 2017). Secondly, some disturbance events, such as fires, droughts or floods, are stochastic in time (duration, intensity and frequency) and space (scale and extent) (Lake 2000; Levin 1992) and their effect on vegetation can be complicated (Pausas and Austin 2001). A static surrogate such as land use cannot capture the spatial and temporal variation in disturbance (Drielsma and Ferrier 2006). By singling out different growth form groups, we implicitly acknowledge that different disturbances influence plant groups in different ways. Advances in data availability (such as routinely updated land use information, or remotely sensed data, including LiDAR, IKONOS, Quickbird, Landsat ETM and Sentinel-2) may offer opportunities to dynamically and iteratively update and refine variables (Leitão and Santos 2019), such as land use, to better predict structure and composition of vegetation at a regional scale. At a global scale, predictors that relate to land use are often inferred from remotely sensed land cover data (see Socioeconomic Data and Applications Center http://sedac.ciesin.columbia.edu).

### Simple and transparent maps of model uncertainty in specific locations

One of the substantial benefits of our modelling approach is the mapped standardised residual error. When displayed as a map, the ‘known error’ at every site is used to infer the ‘unknown error’ across the whole landscape. Here we have used inverse distance weighting to spatially interpolate the residual error between the observed input data and the surrounding areas where errors are unknown. Other approaches, including Kriging, are equally useful (Sajid et al. 2013). The spatial patterns in standardised residual error show that some vegetation indicators are likely to be temporally dynamic, such as forb cover and richness, which have a greater range in the residual error. Whereas vegetation indicators that are relatively stable through time, such as tree richness, show a narrower range in the residual error.

Predictive models are prone to uncertainty. The sources of model uncertainty can arise from any (or all) of the steps outlined in Figure 2, including errors and temporal variation in the response data; inaccurate, imprecise or absent predictor surfaces; or the model may fail to adequately associate (learn) the patterns and relationships between the response data and the predictor variables. Some model uncertainty can be contained by using ensembles of neural network models (Muñoz-Mas et al. 2015) and quantified using sensitivity analyses. In addition to methodological rigor, our analyses and spatial representation of the residual error have provided a simple and transparent means of assessing model uncertainty in specific locations. These maps provide essential information to assist end-users with evaluating the suitability of each model in specific locations and to identify locations for on-ground, visual validation of models.

### Practical applications for mapped vegetation indicators

Quantifying and mapping species composition of vegetation, be that of discrete growth forms or in totality, offers a novel approach for assessing vegetation structure and composition across landscapes. For instance, predictions from these models could be used to inform the habitat preferences for species (e.g. Kissling et al. 2018; Lindenmayer et al. 2018) especially where higher cover or richness of different growth forms contributes to greater habitat complexity (Ashcroft et al. 2017; Brown et al. 1995; Rowe and Speck 2005). These types of biologically-orientated models that can be used to inform habitat-specific occupancy models (McElhinny et al. 2006b), especially the structure and composition of grasses and forbs, which are often overlooked (Gilliam 2007; McElhinny et al. 2005).

In vegetation types where some growth form components are missing yet expected, active restoration can be targeted towards these missing components. For example, Nichols et al. (2010) found that passive regeneration of grasses and forbs was not realised within the ten years post-planting of dominant tree species. We foresee an improved approach to identifying growth forms that could be actively restored to improve the conservation value and enhance some vegetation types. This work has made progress towards targeting restoration efforts to enhance vegetation and habitat quality because we have treated vegetation as a complex of different growth forms.

Additionally, these maps can provide information about vegetation indicators that are useful for estimating the above-ground biomass contribution to carbon accounting or fuel loads which are typically under-detected by remote sensing satellites that collect landscape-scaled imagery, such as Landsat ETM (Lawley et al. 2016), especially the contribution made by non-woody growth forms such as grasses and forbs. Chastain et al. (2006) found that an understorey layer of shrubs made a significant contribution to the carbon content and nitrogen cycling in forests, and importantly, that the substantial understorey layer was not able to be detected using Landsat imagery.

## Conclusions

Often maps of vegetation community type or remotely sensed images of vegetation cover are used to inform conservation and land management. Seldom are the structure and composition of discrete growth forms synthesised from species inventories and mapped as specific and separate indicators. We have focused on advances in three key areas. Firstly, we have identified how response data that are ubiquitous and accessible can be extended to derive community-level structural and compositional vegetation indicators. Secondly, using an extensive suite of predictor data, including environmental and human-induced disturbance surfaces, we used ensemble models to predict and map individual vegetation indicators as 100 m raster layers. We also mapped the interpolated residual error to highlight locations where predictive models have higher uncertainty. These methods can be applied to many regions of the world, especially as new information on the type, frequency and intensity of human-induced and natural disturbance becomes available.

This framework has delivered spatially explicit representations of the structural and compositional indicators of vegetation. We anticipate that these models of vegetation indicators will enhance existing vegetation maps, fill an information gap where maps are not currently complete or consistent and strengthen biodiversity conservation and land management decision-support across a range of applications.

## Authors’ contributions

- conceived the ideas and designed methodology – MM, IO, SF, GN, PGrif, MW, PGib
- extracted response data – MS, MM
- extracted predictor data - MM
- designed software tools – GM
- analysed and interpreted the data – MM, PGrif, TK
- prepared all figures, tables, maps, open data access – MM
- led the writing of the manuscript – MM
- contributed critically to drafting and revising the manuscript – MM, IO, SF, GN, PGrif, MW, PGib, TK
- final approval for publication – ALL

## Acknowledgments

This work was undertaken through collaborative partnerships between NSW Office of Environment and Heritage and the former Border Rivers – Gwydir and Namoi Catchment Management Authorities. Jillian Thonell, Geoff Horn, Simon Smith and Sarah Hill assisted with data preparation. Clive Hilliker advised on the design of Figure 2. Statsoft User Support Forum provided valuable support and information on STATISTICA and neural network analyses. We are grateful to Martin Dillon, David McNellie, Vivian Silvey for valuable comments on the manuscript.

## Supplementary Material

This supplementary material provides details behind the conceptual underpinning of selecting predictor variables, model architecture, software tools and mapped indicators with residual errors for the paper McNellie, M.J., Oliver, I., Ferrier, S., Newell, G., Manion, G., Griffioen, P., White, M., Somerville, M., Koen, T. and Gibbons, P. **Extending site-based observations to predict the spatial patterns of vegetation structure and composition.**

### Supplementary Material S1 Conceptual underpinning of the environmental and disturbance gradients – the predictor variables

We selected a suite of predictor surfaces based on our ecological understanding of the landscape and conceptual understanding of drivers that influence patterns in vegetation; ‘V’ indicates that the specified predictor surface is known to influence the growth and morphology of vegetation extent or type; ‘M’ indicates surfaces included to inform modification or fragmentation of vegetation. ‘V,M’ indicates that within a whole of landscape context, the predictor informs both extent and type of vegetation as well as modification and fragmentation.

Climate surfaces were generated using MTHCLIM in ANUCLIM v6.1 (Xu and Hutchinson 2013) for the 1921-1995 epoch. A 25 m resolution ‘smoothed’ digital elevation model DEM-S (Gallant et al. 2011) was used as the third independent variable needed to interpolate monthly mean climate values. These variables were selected to express various aspects of the annual regimes of temperature, precipitation, and potential evapotranspiration (Box 1981). After ANUCLIM processing, climate surfaces were imported into ArcGIS v10.1, converted to raster layers (25 m) and projected to Lamberts NSW coordinate system. Sharples et al. (2005) note that digital elevation models with a spatial scale of 1 km resolution are normally sufficient for temperature dependent parameters and a 5 km resolution DEM is normally sufficient for rainfall dependent parameters. Table S1 outlines the predictor surfaces used to represent the environmental and disturbance gradients.

**Table S1.**
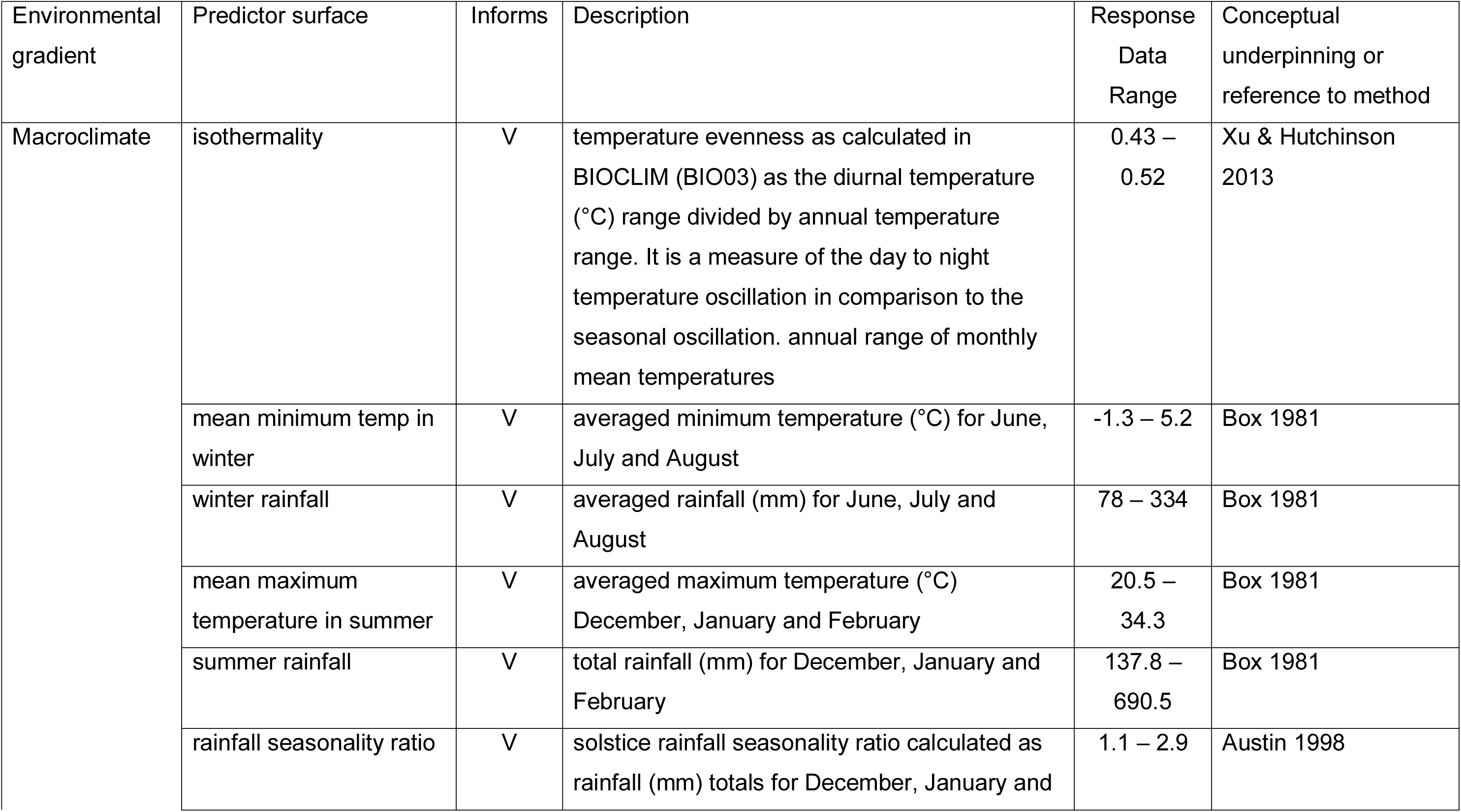
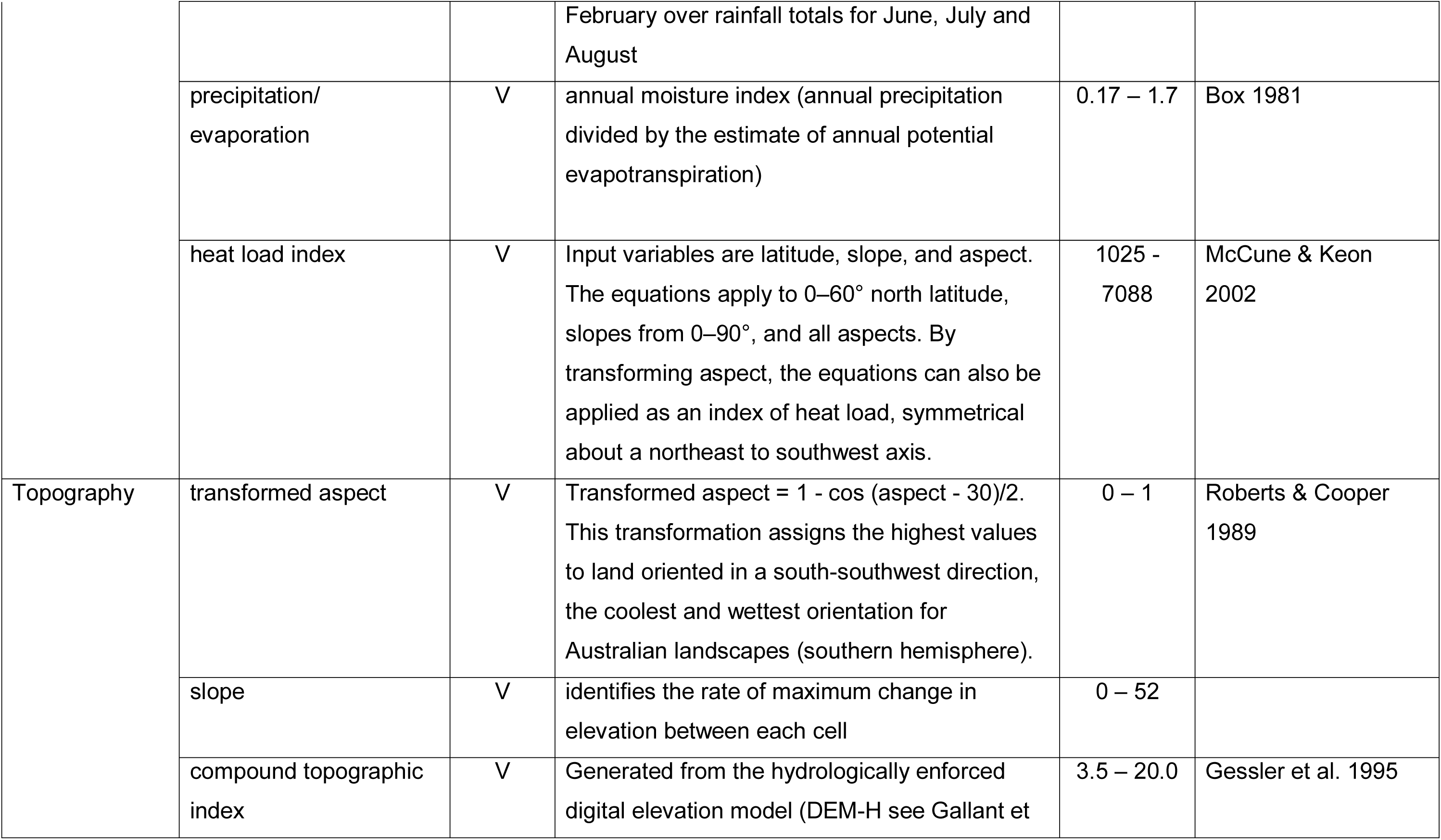
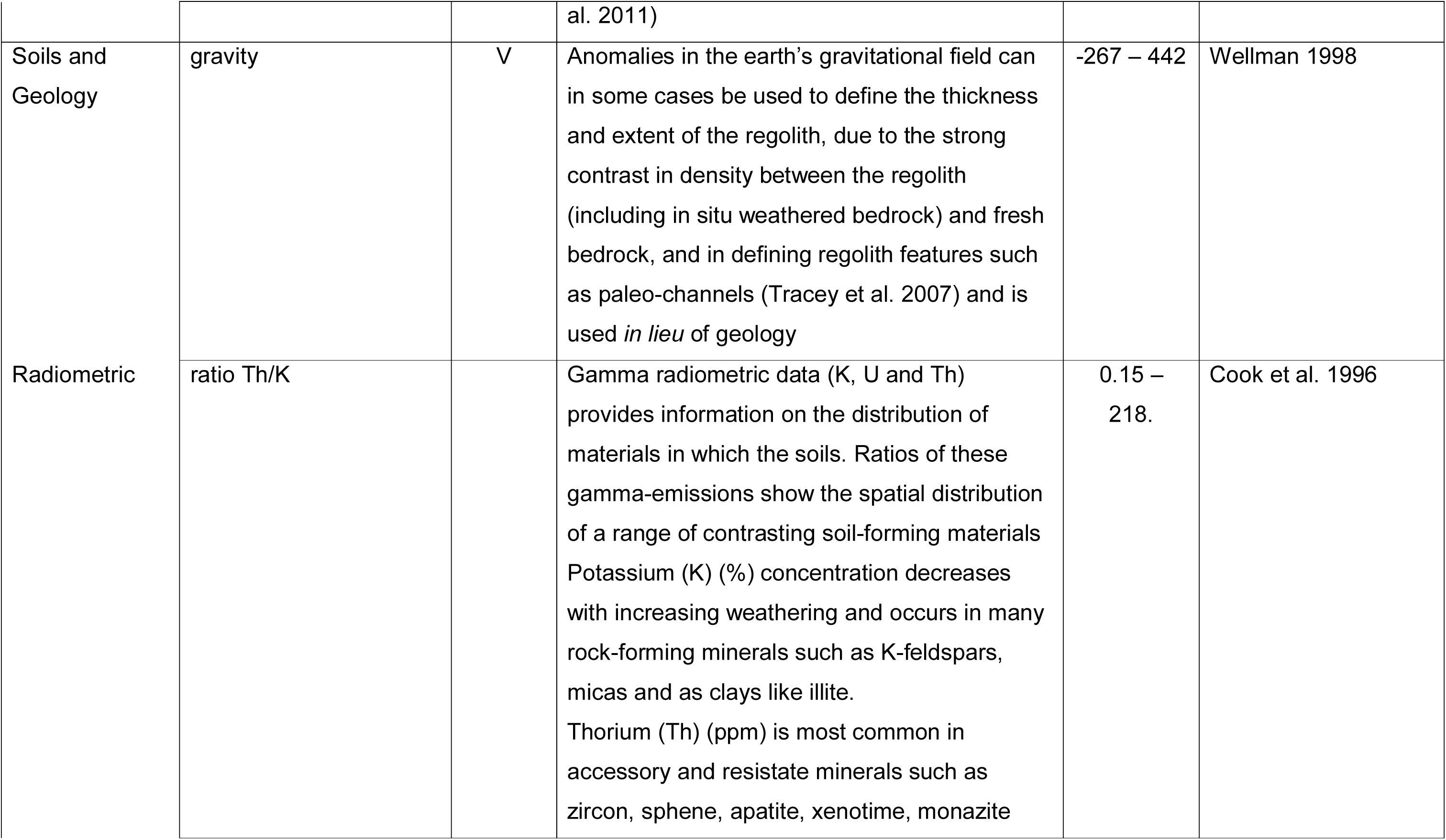
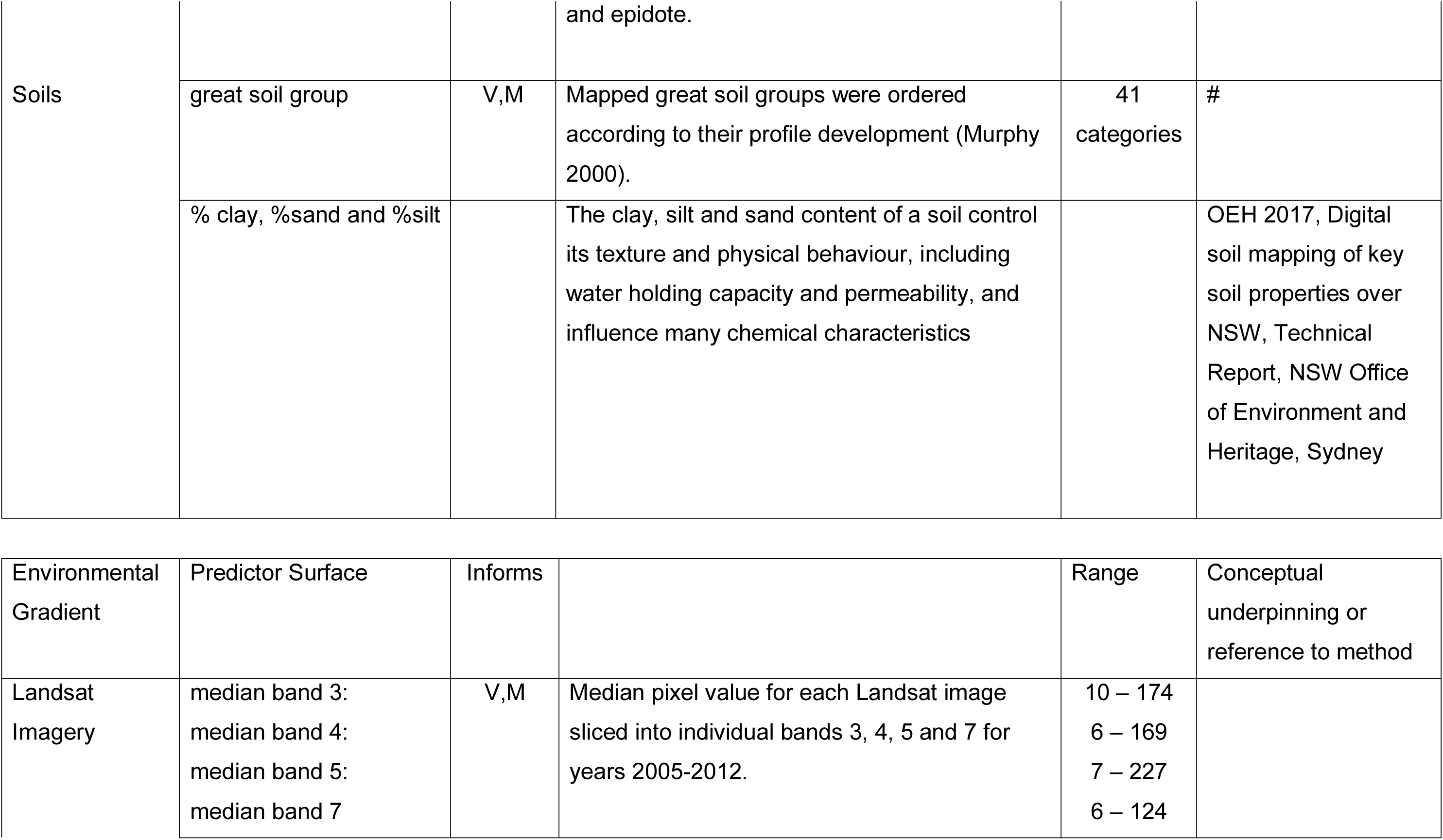
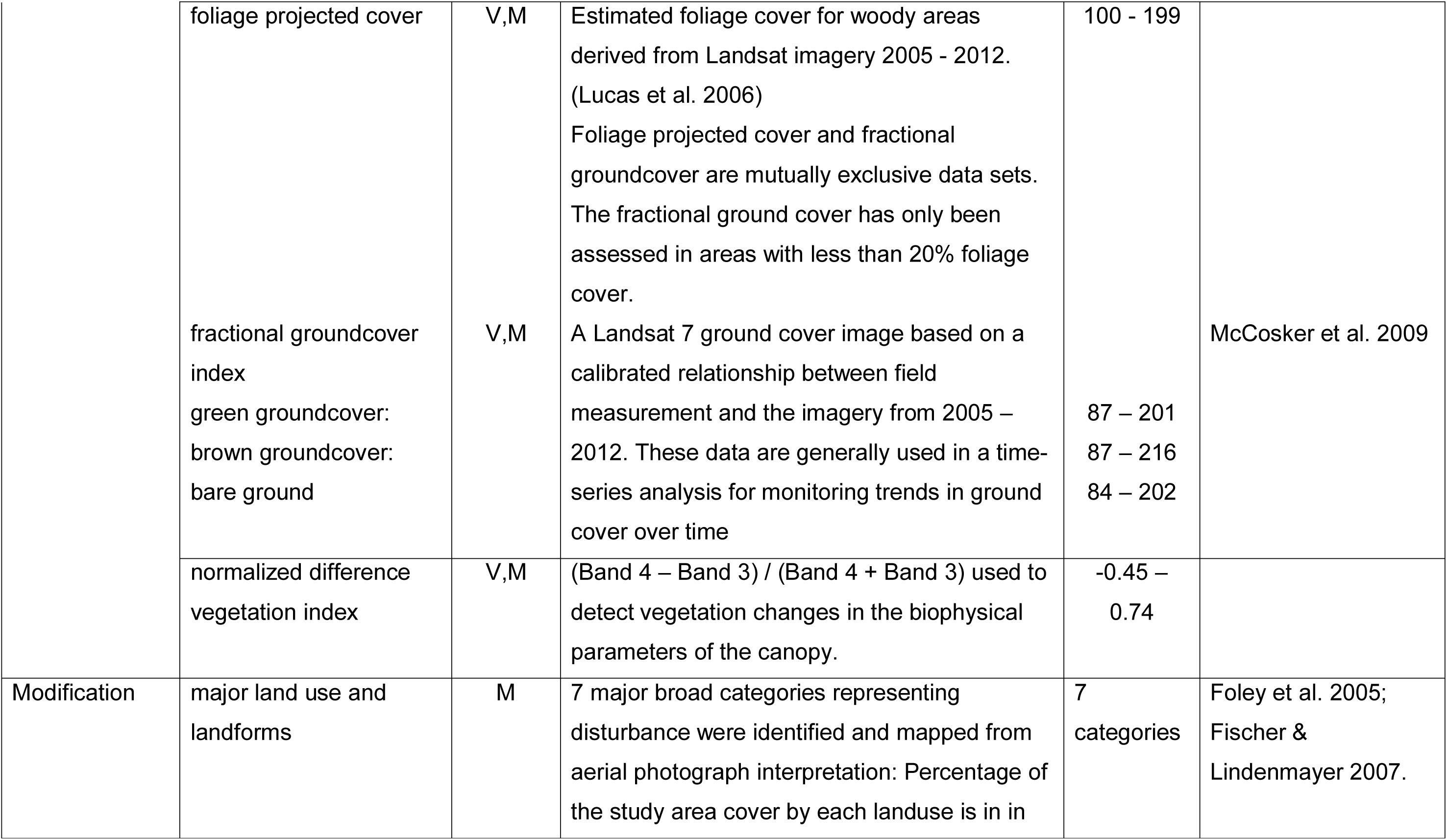
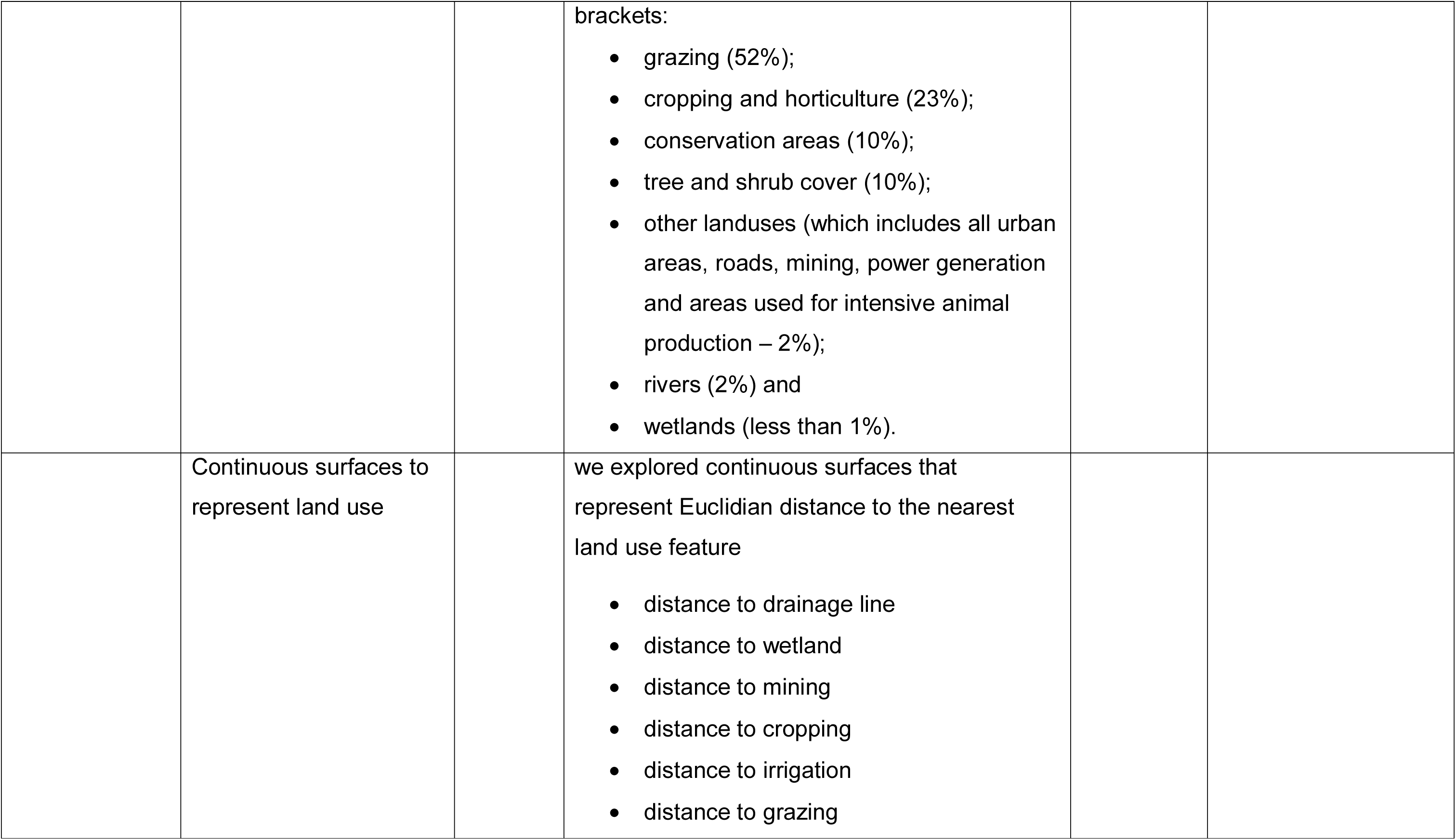
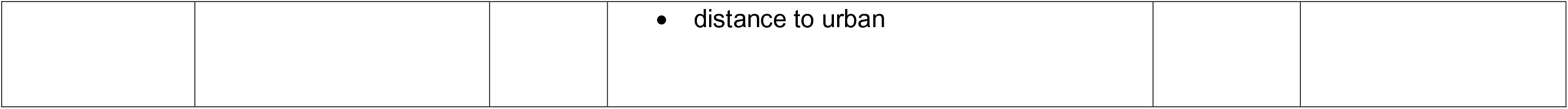
Thirty-four predictor surfaces used in artificial neural network models.

#http://www.environment.nsw.gov.au/Salisapp/resources/spade/metadata/GSG%20Soil%20Types%20of%20NSW%20Metadata%20v131024.pdf

## Supplementary Material S2 Additional information on artificial neural network model architecture

Artificial neural network (ANN) models are used to find patterns in complex data through an iterative ‘train and learn’ process. Training is the process of recognizing the associative patterns and relationships between the response data and environmental predictor surfaces; learning is the process of finding the least amount of error in the patterns and relationships. Training passes information forward through the network via weights and bias; learning informs the network by adjusting weights to reduce the largest errors.

Many types of ANN models exist (see Bishop 1995; Ripley 1996), the most frequent architecture used in ecological applications is a multi-layer perceptron (MLP) feed-forward network with one hidden layer trained by either back-propagation (Rumelhart et al. 1986), or as in our application, the Broyden-Fletcher-Goldfarb-Shanno BFGS algorithm (see Bishop 1995). The MLP framework is used to construct ANN models because their architecture is robust when handling large volumes of complex data.

Below, we discuss the important components of the ANN architecture – observation data, the training algorithm, activation functions and weight decay, predictive output and sensitivity analysis.

### Observation Data

Site locations (X and Y coordinates) were assembled with the intersected point values for each of the 34 environmental and disturbance predictor surfaces (software specifications are outlined in Supplementary Material S3). This matrix formed the response data used to train the ANN. It is generally advised that the dataset contains 10 times more observations than predictor surfaces

### Outlying Observations

Even though ANN models can effectively discount outliers, isolated data points undermine the split-sample hold-out strategy because there are too few cases allocated to either the train, test or validate subsets. This, in turn, has a large effect on the sum-of squares error (Bishop 1995) and consequently, poor predictions can be expected when the validation data contain values outside of the range of those used for training (Maier and Dandy 2000).

### Training algorithm

Training the neural network involves an error algorithm, which finds a set of connection weights that produces an output signal with a small error relative to the observed output. There are many texts that explain the differences between gradient decent and quasi-Newtonian methods, but the fundamental difference is that back-propagation is a gradient decent method and BFGS is a second derivative of the Newtonian method and uses a Hessian error matrix. In terms of applying ANN to ecological modelling, the difference is realised in computational penalties. BFGS require less computational power, which in turn delivers faster models.

### Activation functions

Activation functions introduce nonlinearity into the model. It is the nonlinearity (that is, the capability to represent nonlinear functions) that make ANN models so effective. For both the hidden and output layers we used the negative exponential activation function because the output layers are continuous data, bounded by 0 at the lower with the potential for infinity at the upper end (Bishop 1995).

### Weight decay

To avoid over-fitting the model during training, we used weight decay as an effective form of complex regularisation (Haykin 1994). Minimum weight decay was set to 0.0001 and the maximum to 0.001, which is regarded as weak regularisation. Weight decay trains the network’s sum of squares error function by penalizing larger weights (Statsoft Inc. 2013).

### Predictions

We used Artificial Network Search (ANS) in the Statistica Artificial Neural Network (SANN) module within Statistica v10 software (Statsoft Inc. 2013) to explore multiple configurations of networks. We ran 100 model iterations and varied the number of hidden nodes. We also used weight decay to avoid over-fitting the model during training. For each vegetation attribute, we retained the 25 highest performing models (assessed by the Pearson’s correlation coefficient (*r*)).

The architecture and training parameters used to build the highest 25 performing models are stored as Predictive Model Markup Language (PMML) files. PMML models are a standard format used to represent predictive models across many modelling platforms applications.

### Sensitivity analysis

An indication of the importance of each predictor surface was illuminated using global sensitivity analyses. Sensitivity analyses were calculated after the model was trained and were assessed by comparing the change in the original model error compared to the error obtained by substituting a variable with ‘missing value’. The ratio of ‘original error’ to ‘error when the variable is substituted’, gives an indication of how sensitive the network is to each predictor surface (Statsoft Inc. 2013). Predictor surfaces with sensitivity values less than one cause the model performance to degrade.

## Supplementary Material S3 Software to create data input tables, prediction output tables and manage multiple tables

Creating the training matrix and the prediction matrix and interpreting the prediction output into a raster map was accomplished with the following suite of software tools. There are 32-bit and 64-bit versions these tools. These are all developed under Microsoft.NET and require Microsoft.NET application Framework version 4.0.

- NNExtractFromBinaries.exe (see manuscript Figure 2 Step 2.2 and Step 4.2)
- NNToolsExtractor.exe (see manuscript Figure 2 Step 4.1)
- NNToolsPrediction.exe (see manuscript Figure 2 Step 5.1)

### NNExtractFromBinaries.exe

To build the training matrix, for each of the response data (points), the underlying values from each of the predictor (raster) surfaces were extracted using the NNExtractFromBinaries.exe application (Figure 2 Step 2.2). A comma delimited text file, formatted as Site_ID, X_Coordinate, and Y_Coordinate was used as input into the NNExtractFromBinaries.exe application. The output table was also a comma delimited table with the first three columns formatted as the input table and the comma delimited values for each of the predictor surfaces in subsequent columns. The same tool was used to build the prediction matrix (Figure 2 Step 4.2), using the cell centroids from each of the 100 m grid cells in the study area and the same suite of predictor surfaces.

The predictor surfaces must all be in ESRI Binary Export Format (.flt/.hdr) and can be loaded into the listView in the lower half of the application. The positions of each predictor (order of extraction) can be modified via the up/down arrow buttons and predictors can be eliminated from the listView with the X button. Press Run to start the process of extraction. The status bar will indicate the progress. The csv table is supplied as the input file to train the ANN models.

### NNToolsExtractor.exe

The extent of the study area was prepared as a 100 m raster grid in ESRI Binary Export Format. The centroid of each grid cell in the spatial extent becomes a new site for which the trained network makes a prediction (Figure 2 Step 4.1). These cell centroids are extracted to create the first section of the prediction matrix. The NNToolsExtractor.exe application created a table with the following Format _ID, X_Coordinate, Y_Coordinate where the X and Y coordinates of all the centroids for each of the 11.5 million cells in the study area. Once the X Y locations for every new grid cell are nominated, this table can then be submitted to the NExtractFromBinaries.exe application to extract the predictor data for each record (Figure 2 Step 4.2).

### NNToolsPrediction.exe

Once the predictions have been acquired via the PMML output from the model, the resulting table can be used as input to the NNToolsPrediction.exe application (Figure 2 Step 5.1). The user can select the X and Y fields and importantly, the prediction field and by using the mask grid as a domain will create an ESRI Binary Export grid that will have each valid data cell coded with the predicted value from the ANN model.

## Supplementary Material S4 Comparison of results to assess ensemble model performance when using land use as a categorical or as several continuous layers

Total number of observations (n) for cover models n = 3 481 (with training subset n = 1 741 observations; test subset n = 696 and validation subset n = 1 044) and total number of observations for richness models n = 11 132 (with training subset n = 5 567; test subset n = 2 226 and validation subset n = 3 339). *r* = Pearson’s correlation coefficient; SoS = sum of squares error.

| Indicator | Subset | Land use - continuous |  | Land use - categorical |  |
| --- | --- | --- | --- | --- | --- |
|  |  | *r* | SoS | *r* | SoS |
| Total native cover | Train | 0.83 | 475.44 | 0.85 | 435.86 |
|  | Test | 0.84 | 467.52 | 0.87 | 402.40 |
|  | Validate | **0.79** | 550.88 | **0.83** | 462.91 |
| Tree cover | Train | 0.79 | 120.29 | 0.81 | 114.25 |
|  | Test | 0.79 | 134.69 | 0.80 | 129.94 |
|  | Validate | **0.78** | 131.26 | **0.79** | 125.60 |
| Shrub cover | Train | 0.59 | 136.64 | 0.60 | 134.83 |
|  | Test | 0.56 | 134.22 | 0.57 | 133.04 |
|  | Validate | **0.53** | 136.73 | **0.53** | 136.68 |
| Grass cover | Train | 0.58 | 234.00 | 0.65 | 206.42 |
|  | Test | 0.60 | 218.52 | 0.66 | 191.70 |
|  | Validate | **0.53** | 262.04 | **0.61** | 229.62 |
| Forb cover | Train | 0.36 | 42.41 | 0.41 | 40.39 |
|  | Test | 0.39 | 43.93 | 0.44 | 41.43 |
|  | Validate | **0.36** | 46.78 | **0.42** | 44.31 |
| Total native richness | Train | 0.67 | 83.08 | 0.81 | 51.43 |
|  | Test | 0.68 | 87.12 | 0.83 | 51.81 |
|  | Validate | **0.66** | 89.99 | **0.81** | 53.22 |
| Tree richness | Train | 0.63 | 1.24 | 0.75 | 0.92 |
|  | Test | 0.63 | 1.31 | 0.77 | 0.93 |
|  | Validate | **0.63** | 1.21 | **0.76** | 0.84 |
| Shrub richness | Train | 0.64 | 6.26 | 0.73 | 4.89 |
|  | Test | 0.65 | 6.22 | 0.75 | 4.70 |
|  | Validate | **0.65** | 6.34 | **0.75** | 4.82 |
| Grass richness | Train | 0.53 | 9.92 | 0.70 | 6.99 |
|  | Test | 0.54 | 10.26 | 0.72 | 6.88 |
|  | Validate | **0.52** | 10.14 | **0.72** | 6.72 |
| Forb richness | Train | 0.54 | 18.71 | 0.69 | 13.57 |
|  | Test | 0.56 | 18.43 | 0.72 | 12.95 |
|  | Validate | **0.53** | 18.86 | **0.70** | 13.30 |

## Supplementary Material S5 The spatially explicit maps and accompanying residual maps for each indicator

**Figures S1-S5** Maps of predicted vegetation indicator and their standardised residual error. The standardised residual error between predicted and observed values was calculated for each prediction at the site and interpolated across the landscape using inverse distance weighting (IDW) in ArcGIS Spatial Analyst. These values represent areas where the model has under estimated (positive values) or over estimated (negative values). Residual error maps were classified so that residual error (−1 and 1) are white; increasing residual error (between 1 to 2 and −1 to −2) as shaded lighter, and areas where there is higher uncertainty (less than −3 or greater than 3) are darker.

**Figure S1.**
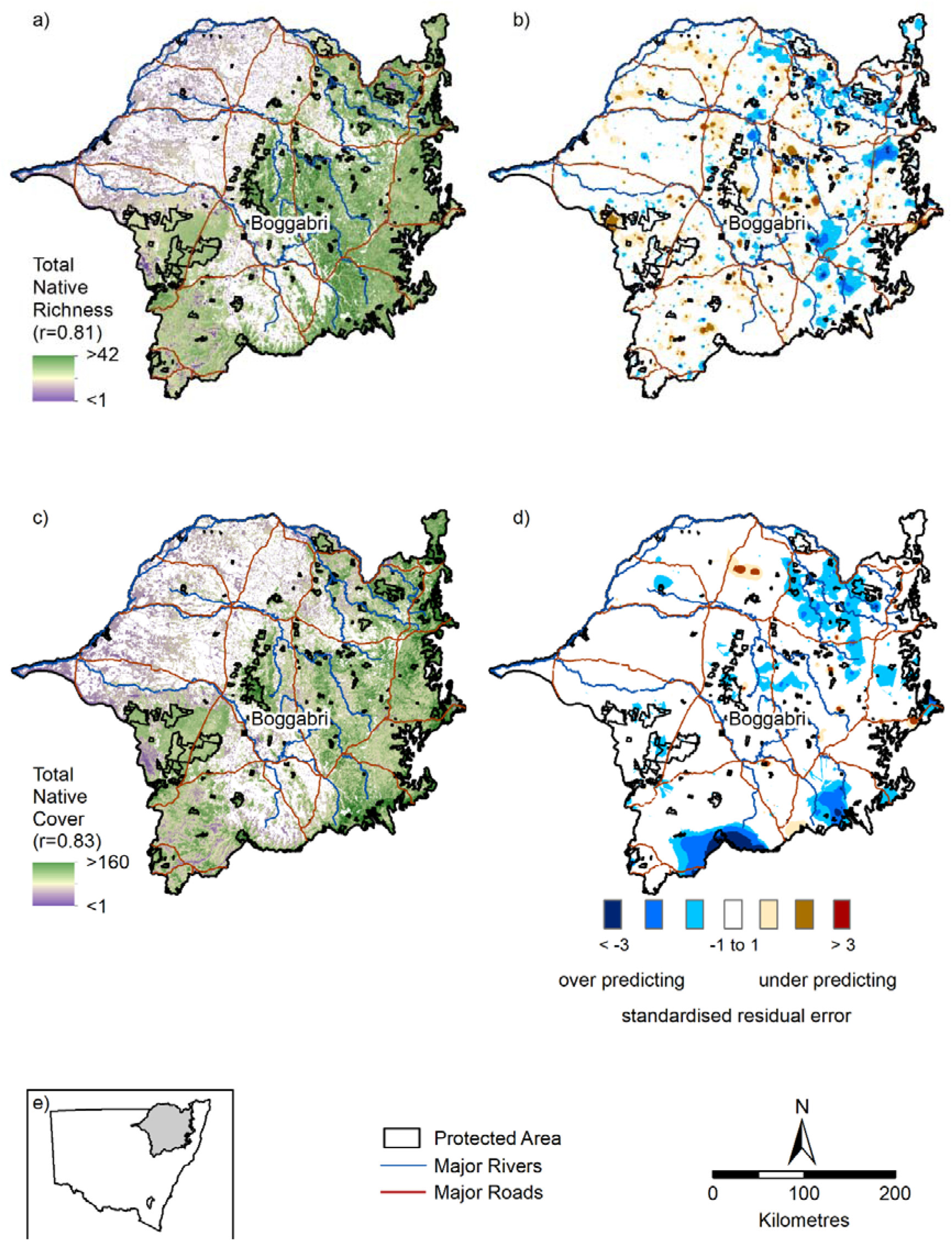
a) total native richness; b) standardised residual error for total native richness; c) total native cover (%); d) standardised residual error for total native cover and e) inset map showing study area

**Figure S2.**
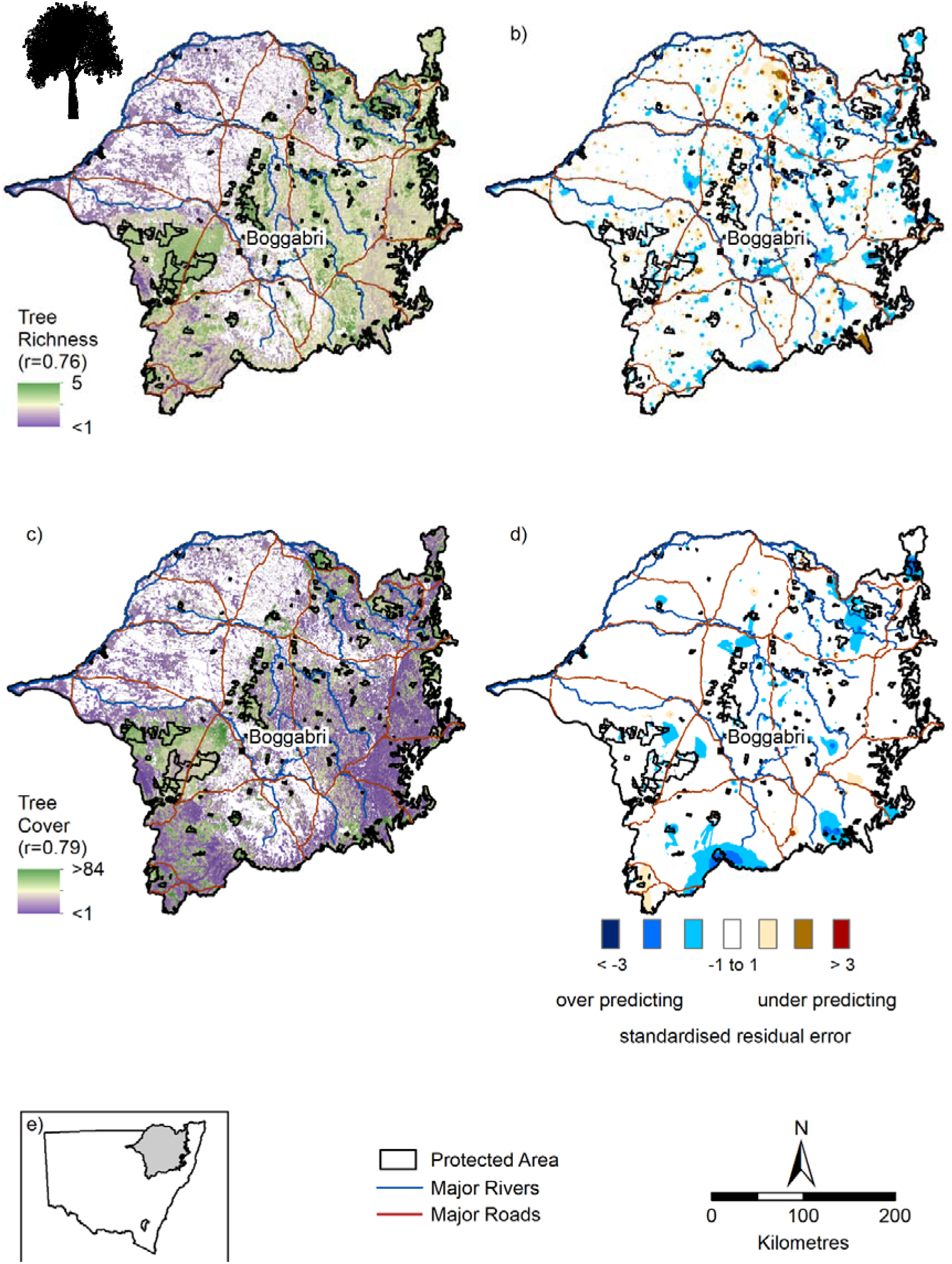
a) native tree richness; b) standardised residual error for native tree richness; c) native tree cover (%); d) standardised residual error for native tree cover; and e) inset map showing study area

**Figure S3.**
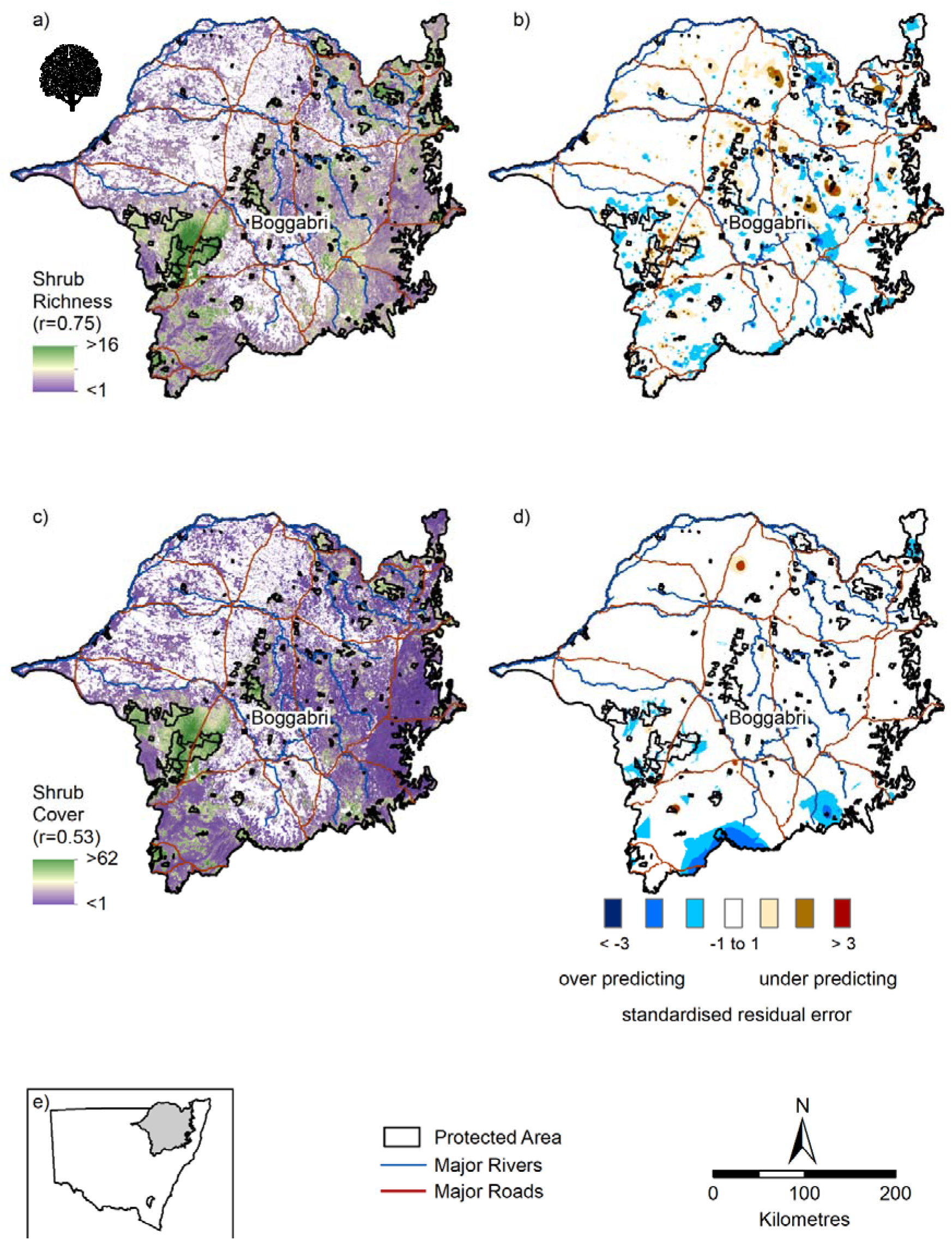
a) native shrub richness; b) standardised residual error for native shrub richness; c) native shrub cover (%); d) standardised residual error for native shrub cover and e) inset map showing study area

**Figure S4.**
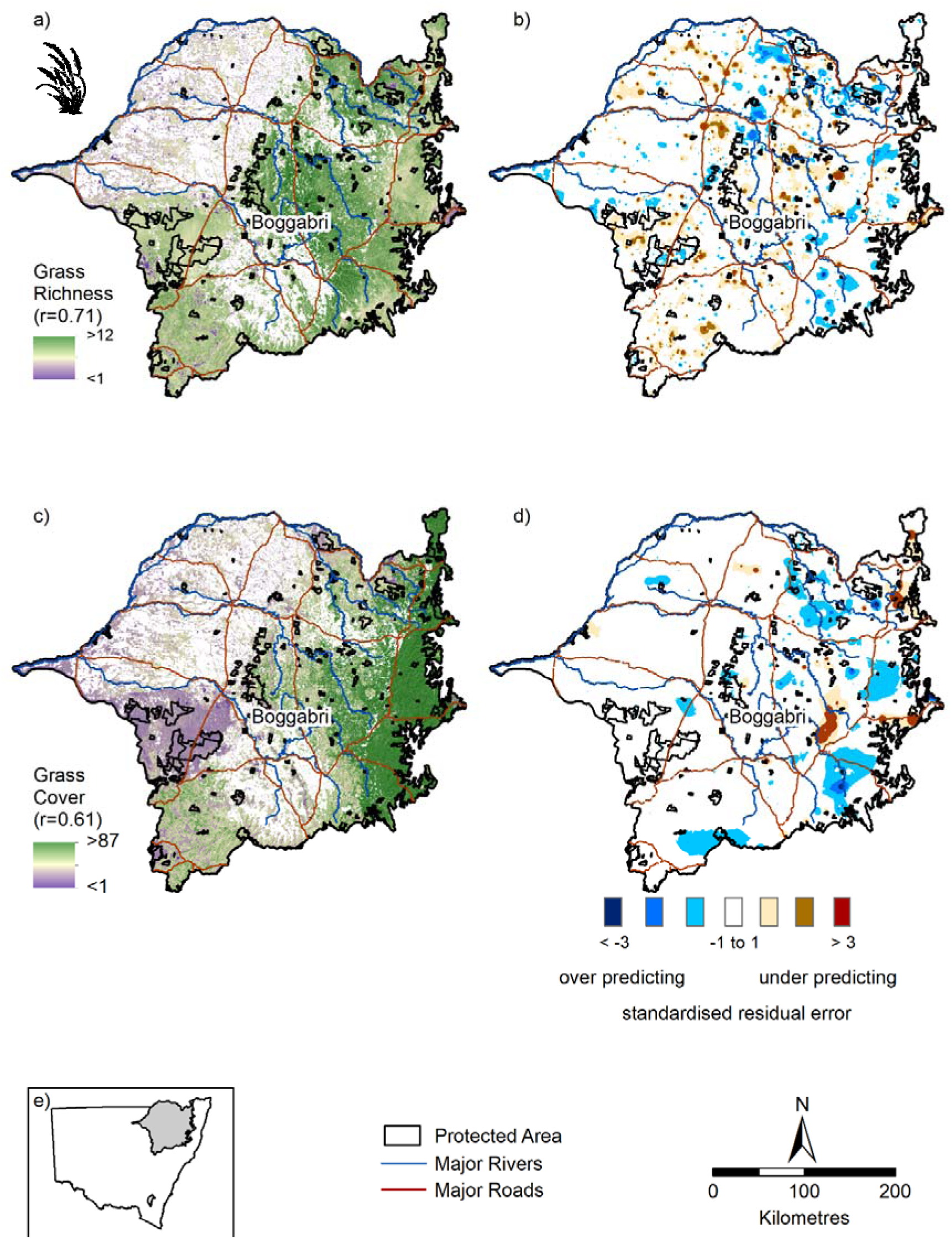
a) native grass richness; b) standardised residual error for native grass richness; c) native grass cover (%); d) standardised residual error for native grass cover and e) inset map showing study area

**Figure S5.**
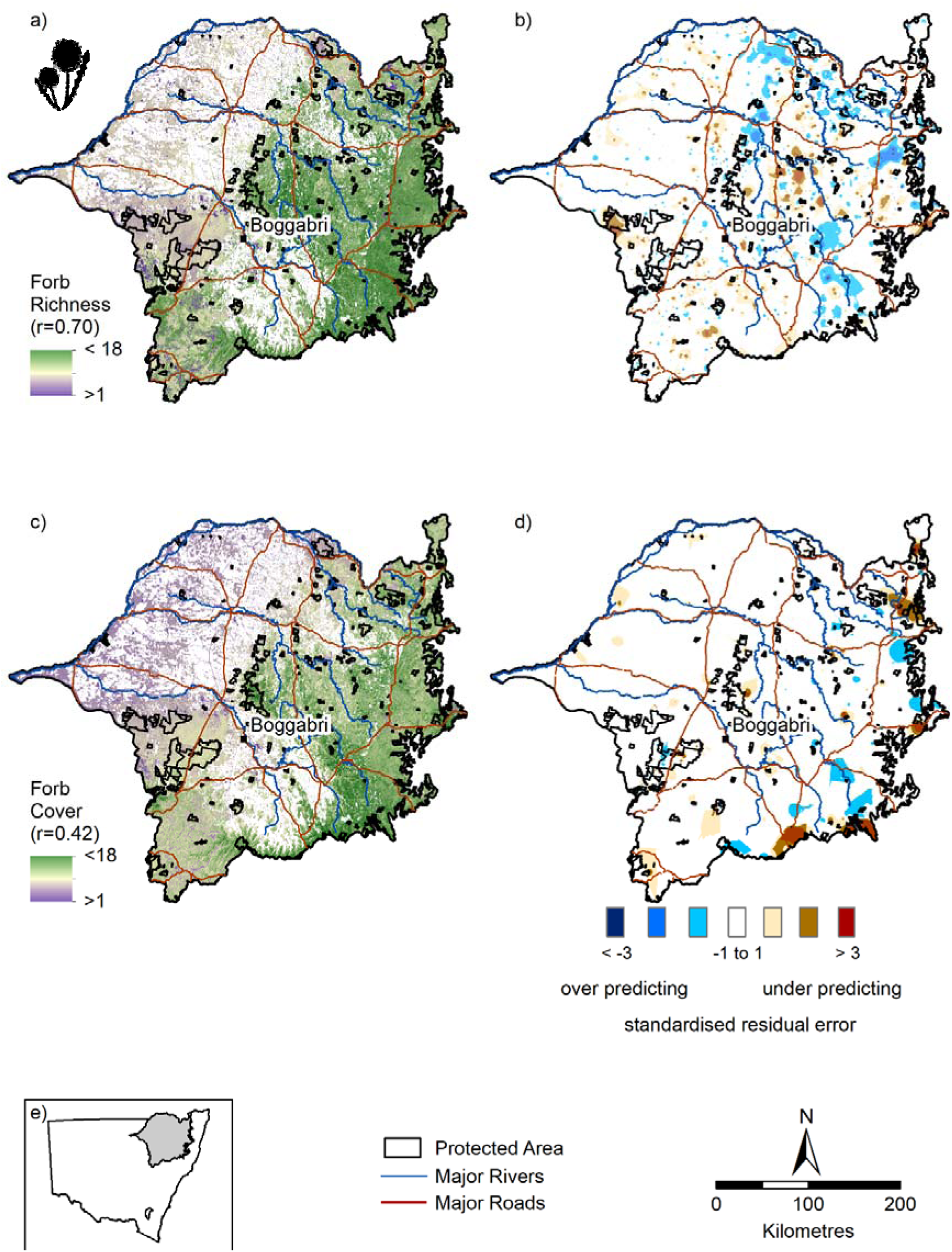
a) native forb richness; b) standardised residual error for native forb richness; c) native forb cover (%); d) standardised residual error for native forb cover and e) inset map showing study area

